# An auto-inhibited state of protein kinase G and implications for selective activation

**DOI:** 10.1101/2022.04.28.489861

**Authors:** Rajesh Sharma, Jeong Joo Kim, Liying Qin, Philipp Henning, Madoka Akimoto, Bryan VanSchouwen, Gundeep Kaur, Banumathi Sankaran, Kevin R. MacKenzie, Giuseppe Melacini, Darren E. Casteel, Friedrich W. Herberg, Choel Kim

## Abstract

Cyclic GMP-dependent protein kinases (PKGs) are key mediators of the nitric oxide/cGMP signaling pathway that regulates biological functions as diverse as smooth muscle contraction, cardiac function, and axon guidance. Campaigns targeting nitric oxide synthases and cyclic nucleotide phosphodiesterases in this signaling axis suggest that understanding how cGMP differentially triggers mammalian PKG isoforms could lead to new therapeutics that inhibit or activate PKGs. Alternate splicing of PRKG1 transcripts confers distinct leucine zippers, linkers, and auto-inhibitory pseudo-substrate sequences to PKG Iα and Iβ that result in isoform-specific activation properties, but the mechanism of enzyme auto-inhibition and its alleviation by cGMP is still not well understood. Here we present a crystal structure of PKG Iβ in which the auto-inhibitory sequence and the cyclic nucleotide binding domains are bound to the catalytic domain, providing a snapshot of the auto-inhibited state. Specific contacts between the PKG Iβ auto-inhibitory sequence and the enzyme active site help explain isoform-specific activation constants and the effects of phosphorylation in the linker. We also present a crystal structure of a PKG I cyclic nucleotide binding domain with an activating mutation linked to Thoracic Aortic Aneurysms and Dissections. Similarity of this structure to wild type cGMP-bound domains and differences with the auto-inhibited enzyme provide a mechanistic basis for constitutive activation. We show that PKG Iβ auto-inhibition is mediated by contacts within each monomer of the native full-length dimeric protein, and using the available structural and biochemical data we develop a model for the regulation and activation of PKGs.

## Introduction

The second messenger cyclic guanosine monophosphate (cGMP) regulates a myriad of physiological processes including cellular growth, smooth muscle contractility, cardiovascular homeostasis, inflammation, sensory transduction, bone growth, and neuronal plasticity and learning (Battye et al., 2011; Francis et al., 2010; Hammond and Balligand, 2012; Klinger and Kadowitz, 2017). In eukaryotes, cGMP-dependent protein kinases (PKGs) transduce intracellular cGMP levels, which change in response to extracellular signals, into phosphorylation of target proteins that control cell activities. Wildtype PKGs show minimal kinase activity in the apo state but are strongly, cooperatively, and selectively activated by sub-micromolar concentrations of cGMP. The regulation and mechanism of action of PKGs is of fundamental importance to understanding the basis of biological responses to cyclic nucleotides and could reveal therapeutic strategies that would complement current approaches to colon cancer, hypertensive heart disease, pulmonary hypertension, osteoporosis, and chronic pain (Browning et al., 2010; Feil et al., 2003; Kalyanaraman et al., 2018; Klinger and Kadowitz, 2017; Luo et al., 2014).

PKGs belong to the AGC family of protein kinases and have N-terminal regulatory (R) and C-terminal catalytic (C) domains. The PKG Iα and Iβ isoforms are splice variants with different N-terminal regions of 89 or 104 residues but the same C-terminal 582 residues that form cyclic nucleotide binding (CNB) and catalytic domains (Francis and Corbin, 1994; Hofmann et al., 2009). PKG I induces smooth muscle relaxation by lowering intracellular calcium or activating myosin phosphatase (Schlossmann et al., 2000; Surks et al., 1999). PKG II is produced from a different gene with 63 % identity and regulates ERK activation required for bone growth, and trafficking of cystic fibrosis transmembrane conductance regulator (Pfeifer et al., 1996; Rangaswami et al., 2009; Serulle et al., 2007; Vaandrager et al., 1998). Differences between the isoforms are important to their function (Schlossmann and Hofmann, 2005) and may be exploitable for selective targeting by small molecules.

Much of our understanding of the mechanism of action of PKGs derives from studies of isolated domains. The R-domain comprises a leucine zipper (LZ) domain, a linker region with an auto-inhibitory (AI) sequence, and two cyclic nucleotide binding (CNB-A and -B) domains (***Figure 1A***). All three PKG isoforms have unique LZ and AI domains. The LZ domains mediate homodimerization and recruit isoform-specific interacting proteins (Casteel et al., 2010; Qin et al., 2015; Reger et al., 2014). The isoform-specific AI sequences contain pseudo-substrate motifs with the target S/T replaced by <u>G</u> or <u>A</u> (RAQGIS in PKG Iα, <u>K</u>RQAIS in PKG Iβ, and AKA<u>G</u>VS in PKG II) that can bind the catalytic cleft of the C-domain, blocking substrate access (Francis et al., 1996). The CNB-A and -B domains bind cGMP with differing affinities and selectivities (Huang et al., 2014b; Kim et al., 2011) and are connected by interdomain helices. Each CNB includes a β subdomain flanked by helical subdomains (Berman et al., 2005; Rehmann et al., 2007). Cyclic nucleotide pockets comprise the phosphate binding cassette (PBC), base binding region (BBR), and capping residue (Das et al., 2009). The side chain of a PBC arginine that is conserved in all CNBs contacts a non-bridging oxygen in the cyclic phosphate. A mutation that causes familial Thoracic Aortic Aneurysms and Dissections (TAAD) maps to this arginine of CNB-A (PKG Iα residue 177), and the mutation to glutamine constitutively activates the enzyme (Guo et al., 2013). In PKG I, CNB-B includes a BBR arginine that specifically recognizes cGMP over cAMP; CNB-A lacks this arginine and binds either cyclic nucleotide with similarly high affinity (Huang et al., 2014b; Kim et al., 2011). Capping residue Y351 at the C-terminal loop of PKG Iβ CNB-B shields the guanine moiety from solvent and is therefore also referred to as a lid (Huang et al., 2014b).

**Figure 1.**
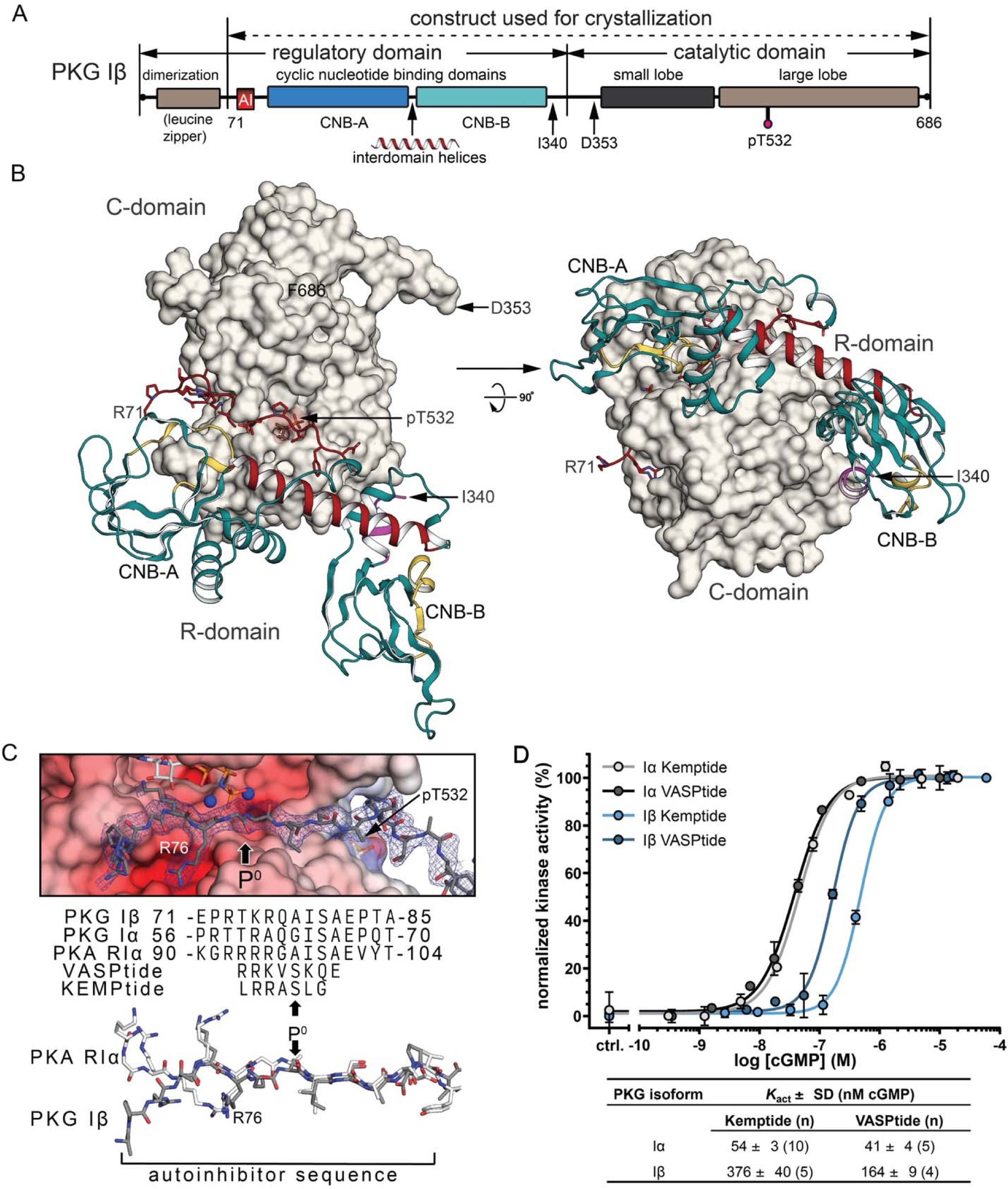
Overall structure of the R:C holoenzyme complex. (A) Domain organization of PKG Iβ and the construct used for crystallization. AI, auto-inhibitory sequence; CNB, cyclic nucleotide binding domain. (B) Overall structure of the PKG Iβ R:C complex (71-686). The R-domain is shown in cartoons while the C-domain is shown in grey surface. CNB-A and B are colored in teal, PBC in yellow, αA-helix in magenta, and the interdomain helices in red. AI and phosphorylated pT532 are colored in red sticks. All structure images were generated using PyMOL (Delano -Scientific). (C) AI docking to the active site. Top: AI is shown with electron density (2*F_o_-F_c_* at σ=1.0). The phosphorylation site (P^0^) in models and sequences is marked with arrows. Electrostatic potential surface is shown for the C-domain active site. Bottom: Alignment of AI sequences of PKG Iβ, Iα, and PKA RIα with substrates VASPtide and Kemptide. (D) Isozyme differences in cGMP-dependent activation of PKG Iα (grey) and Iβ (blue) are revealed by activity measurements. Activation constants (*K*_act_) also differed with varying substrates. Both isoforms require less cGMP for half-maximal activation when using VASPtide (dark grey and dark blue) compared to Kemptide (light grey and light blue). Data points show the mean of duplicates with error bars indication the standard deviation (SD). *K*_act_ values are given as mean of *n* measurements ± SD. Additional data related to these fits are presented in ***Figure supplement 3***.

By analogy to PKA, the PKG R-domain is thought to bind via its AI sequence to the catalytic C-domain active site to form an auto-inhibitory R:C state; binding of cGMP to the R-domain destabilizes this state and allows active kinase to phosphorylate downstream substrates. In contrast to PKG, activation of the auto-inhibitory PKA R:C state by cAMP dissociates the auto-inhibited tetramer into separate R and C subunits. In previous work, we showed that a PKG I R-domain comprising two CNB domains but lacking the dimerization domain makes R:R intermonomer contacts through the cGMP bound at CNB-A and at a kink in the interdomain linker between the two CNBs (Kim et al., 2016). Disrupting this interface with point mutations increases the activation constant in dimeric full-length PKG Iβ, suggesting a role for the domain:domain contacts that mediate this interface in cooperatively stabilizing the activated state of the holoenzyme (Kim et al., 2016) despite the auto-inhibitory sequences tethered in proximity to the C-domains.

Here we present the first crystal structure of PKG Iβ in which the regulatory domain is free of cyclic nucleotide and adopts an auto-inhibited state bound to the catalytic domain. The R:C interface reveals specific contacts between the PKG Iβ AI sequence and the active site that help to explain isoform-specific activation constants and the activating effects of phosphorylation in the linker. Comparison with the structures of isolated PKG R- and C-domains shows that the R-domain undergoes local conformational changes and domain rearrangements to assemble a surface that contacts the catalytic core of the C-domain and inhibits its activity. We also present a high resolution structure of the CNB-A domain with the TAAD mutation, which reveals a closed cGMP pocket. The similarity of this mutant domain to wild type cGMP-bound structures and its differences with the open pocket in the auto-inhibited enzyme explain the constitutive activation of the holoenzyme in TAAD. We show that PKG Iβ auto-inhibition is mediated by contacts within each monomer of the native full-length dimeric protein, and using the available structural and biochemical data we develop a model for the regulation and activation of PKGs.

## Results

### Overall structure

The asymmetric unit of our crystals of PKG Iβ 71-686 contains two PKG Iβ peptide chains (see ***Figure 1A*** for domain structure) without cGMP (***Table supplement 1***). Structures of the individual domains are very similar to those of previously reported isolated domains, validating these (Huang et al., 2014b; Kim et al., 2011; Qin et al., 2018) and providing opportunities to infer the effects of inter-domain interactions. Stretches of ∼15 amino acids connecting the R- and C-domains lack electron density in both chains (the R-domain traces end at residue 341 or 342, and the C-domain traces begin at residue 353 or 355), so which R-domain is connected to which C-domain within the crystal is undetermined. The disordered regions for which the crystal has no density are helical in structures of the isolated CNB-B domain and of the catalytic domain (***Figure supplement 1***), but our recent experiments with full-length PKG Iβ show a high rate of backbone amide hydrogen/deuterium exchange in this region, whether PKG Iβ is auto-inhibited or activated (Chan et al., 2020). The shortest connecting path in the crystal (about 15 Å) could be spanned by a kinked helix to give a dimer inhibited in trans, with the R-domain of one monomer bound to the C-domain of the other (***Figure 1B***). In solution, however, this construct is monomeric, and its experimental radius of gyration from small angle X-ray scattering is in good agreement with the calculated radius of gyration for a monomer formed by the alternate topology (***Table supplement 2***).

The entire C-domain shows clear density in both chains; besides the missing residues discussed above, only the first 2 residues at the N-terminus of both chains (residues 71-72) and five residues that follow the AI sequence (residues 85-89) in one chain are missing. The R-domain AI sequence occupies the C-domain active site (***Figure 1C***), and the contacts in this region partly explain the differences in cGMP activation between PKG Iα and Iβ (***Figure 1D***). The C-domain with AMP-PNP:Mn^2+^ bound at the ATP site shows a ‘closed’ conformation similar to our previous structure of the isolated C-domain with an active site inhibitor (Qin et al., 2018). Both CNB-A and CNB-B PBCs adopt ‘open’ conformations with respect to the β-subdomains, similar to previous structures for CNBs without cyclic nucleotide (Boettcher et al., 2011; Kim et al., 2007; Lu et al., 2019). Despite these gross similarities, local conformational changes enable the overall architecture of the R-domain in the R:C complex to differ substantially from previous R-domain structures, as discussed below.

### The PKG Iβ R:C interface

We describe the contacts between the PKG Iβ R and C-domains in terms of four docking subsites composed of R- and C-domain structural elements (as shown in ***Figure 2A***) whose total buried interface area is ∼2400 Å^2^ (Krissinel and Henrick, 2007). The R-domain AI sequence occupies the C-domain active site (***Figures 1C and site 1 in 2B***). The helical subdomain of CNB-A, including PBC-A and N3A^A^, combines with the AI to form an extended surface that binds the αG-helix of the C-domain (names for catalytic domain elements follow the established convention for kinases (Johnson et al., 2001), and names for CNB elements follow the convention from (Berman et al., 2005; Kornev et al., 2008)). At the third site, the interdomain helices and the αA helix of CNB-B shield the activation loop of the C-domain, and at the fourth, the αB helix of CNB-B docks to an S-shaped loop (residues 610-625) between the αH and αI helices at the bottom of the C-domain (***Figure 2B***). In the R:C complex we present here, the interdomain linker between the CNB domains forms a single helix that allows the R-domain to clamp the C-domain between its two CNB domains.

**Figure 2.**
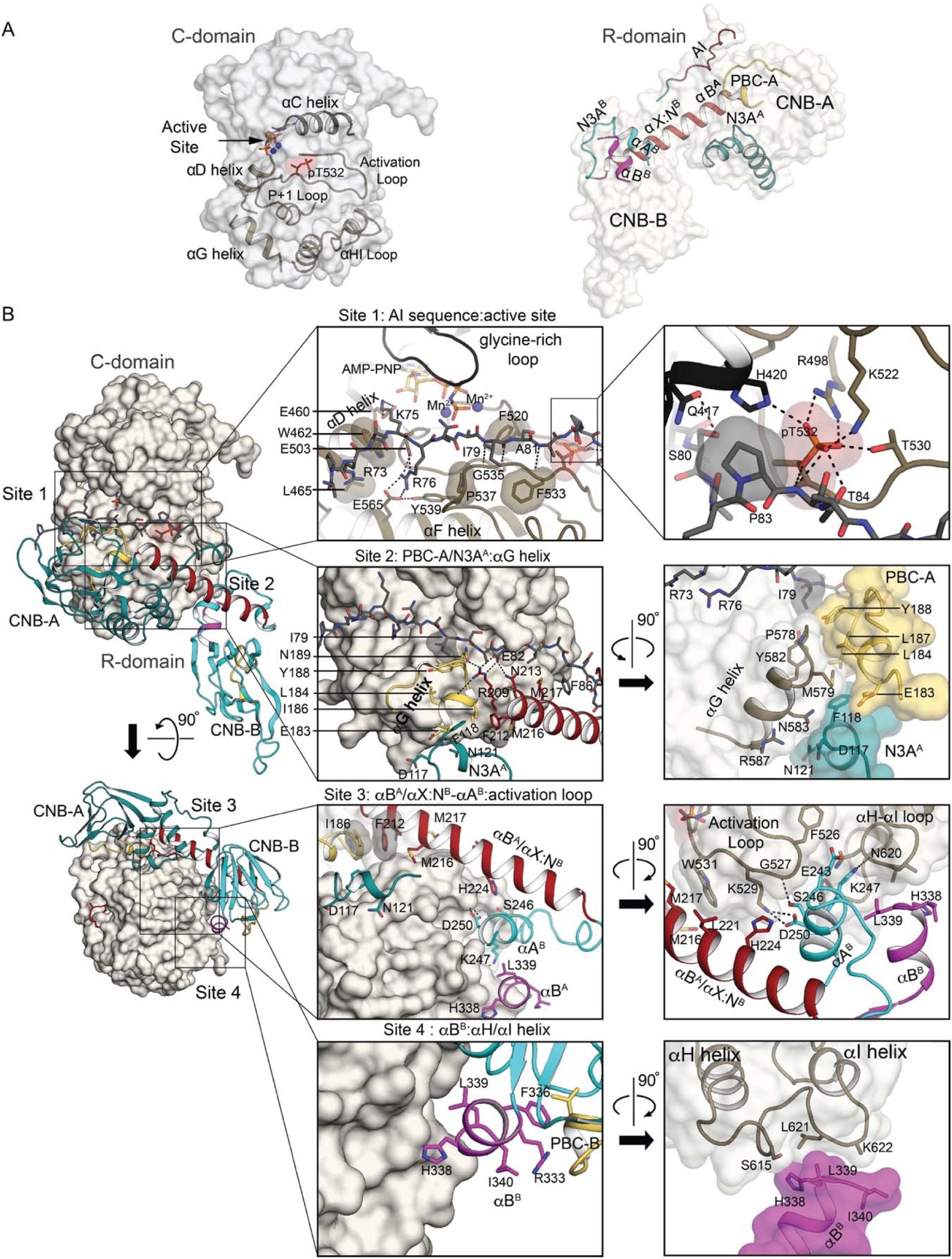
Interactions at the R:C interface. (A) Key structural elements that form the R:C interface. The C and R domains (left and right) are displayed in isolation as transparent surface with key elements labeled and shown as cartoons using the same color scheme as in *Figure 1B*. (B) Detailed R:C interactions with the C-domain presented as a surface and the R-domain portrayed in cartoon using the same color scheme. Left two panels: Two orthogonal views of the auto-inhibited complex with outlines identifying the regions discussed in the text as interaction sites 1 through 4. Middle four panels: Zoomed-in views of the four interaction sites with key residues labeled and shown as sticks, and hydrogen bonds shown as dotted lines. Right four panels: further zoomed-in or rotated views of the four interaction sites.

At site 1, the AI sequence docks to the active site of the C-domain (***Figures 1C and 2B***) burying a surface area of 730 Å^2^. The final model shows clear electron density for residues 72 to 84 in both chains, though no density is visible for the R73 side chain in one of the chains. The AI region docks to the active site cleft through polar and non-polar interactions (right zoom-in panel of Site 1 in ***Figure 2B***). Residues I79, S80 and A81, which are common to PKG Iα and Iβ, form a three residue antiparallel β-sheet with the P+1 loop of the C-domain: the backbone amide and carbonyl oxygen of I79 form hydrogen bonds with the carbonyl oxygen and amide of G535 at the P+1 loop, and the amide of A81 forms a hydrogen bond with the carbonyl oxygen of F533. Residues R73 (P-5), K75 (P-3), and R76 (P-2), which are specific to PKG Iβ, interact with the αD and αF C-domain helices through their side chains. The aliphatic side chains of R73 and K75 sandwich the side chain of W462 at the αD helix of the C-domain, the K75 side chain hydrogen bonds to the D460 side chain at the αD helix, and the R76 side chain forms salt bridges with E503 of the catalytic loop and E565 of the F helix (***Figure 2B***); the importance of the contacts made by these residues to auto-inhibition was first established by activity measurements of proteolytic fragments (Francis et al., 1996). The side chain of the pseudo-substrate P+1 residue, I79, docks to a hydrophobic pocket formed by F533, P537 and Y582 of the C-domain and makes van der Waals contact with the aromatic side chain of Y188 of PBC-A (right zoom-in panel of site 2 in ***Figure 2B***). E82 makes a strong hydrogen bond with R209 of the B helix and positions it to interact with N189 at the helical tip of PBC-A helping to organize site 2 of the R-domain.

In the PKA RIα:Cα complex, four consecutive arginine residues in the RIα AI that interact with acidic residues at the active site of Cα contribute to the high affinity between PKA subunits (Kim et al., 2007). Compared to the PKA RIα:C complex, PKG Iβ has fewer positively charged residues in the AI sequence and shows fewer hydrogen bonds at Site 1 (***Figure 1C***); modeling interactions for PKG Iα suggests that it makes even fewer stabilizing contacts. If the conserved residues I79, A81, and E82 position the PKG Iα AI pseudo-substrate to contact the C-domain as in the PKG Iβ structure, then mapping the PKG Iα P-5, P-3 and P-2 residues T57, R59 and A60 (***Figure 1C***) onto the PKG Iβ Al structure suggests the loss of two salt bridges with the active site at P-2 and of non-polar interactions with the αD helix at P-5. These differences could weaken the R:C interface and may contribute to the lower activation constant and higher basal activity for PKG Iα compared to PKG Iβ (***Figures 1D and Figure supplement 3***).

At site 2, the C-domain αG helix provides a non-polar docking surface that contacts CNB-A at PBC-A and N3A^A^ (site 2 in ***Figure 2B***). A cluster of non-polar residues including M579 and Y582 in the αG helix provide a docking surface for L184, L187 and Y188, which comprise the hydrophobic tip of the CNB-A PBC that adopts an ‘open’ conformation with respect to the β-subdomain (zoom-in panels of site 2 in ***Figure 2B***). The rest of the αG-helix docks with the 3_10_-helix of N3A^A^, for a total buried hydrophobic surface area at this site of 360 Å^2^.

At site 3, the activation loop of the C-domain contacts three elements of the R-domain: a linker segment immediately C-terminal to the AI sequence, the interdomain helix that connects CNB-A and CNB-B, and the αA helix of CNB-B. The total surface area buried at this site is 402 Å^2^. The linker segment interacts with phosphothreonine (pT) 532 and adjacent regions, while the interdomain helix shields the rest of the activation loop. As seen in the right zoom-in panel of site 1 in ***Figure 2B***, P83 and T84 in the linker make non-polar and hydrogen bonding contacts with pT532. P83 stacks with the side chain of H420 (in the C-domain αC helix); the H420 side chain Nε donates a hydrogen bond to the phosphate moiety of pT532 and shields it from solvent. The side chain hydroxyl and backbone amide of T84 donate hydrogen bonds to the phosphate of pT532, which is held in place by two hydrogen bonds from the R498 side chain and one from the T530 side chain. Replacing the analogous PKG Iα phosphorylatable threonine (T517) with alanine results in loss of kinase activity, while replacing it with glutamate retains basal activity, indicating that phosphorylation at this site of the activation loop is required for catalysis (Feil et al., 1995). The contacts that we report here between the R-domain and this key phosphorylated threonine should help stabilize R:C interaction and thus favor auto-inhibition.

The local architecture of this region explains why phosphorylation of S80 activates PKG Iβ (Smith et al., 1996). In our structure, the S80 side chain hydroxyl donates a hydrogen bond to the A81 carbonyl oxygen, receives a hydrogen bond with Q417 of the αC helix, and is just 3.5 Å from the αC-helix H420 side chain that contacts pT532 (right zoom-in panel of site 1 in ***Figure 2B***). In this state, S80 is not accessible for phosphorylation. Phosphorylated S80 would be unable to hydrogen bond to the A81 carbonyl oxygen, and would experience charge repulsion with the phosphorylated activation loop and αC-helix of the C-domain. At the analogous position in PKG Iα, the phospho-mimetic mutation S65D renders the enzyme almost completely constitutively active (Busch et al., 2002); this mutation can be accommodated in our structure only by adjusting the Q417 rotamer away from the AI and pointing the S65D rotamer away from the C-domain. Interestingly, mutations S65A in PKG Iα and S80A in PKG Iβ both raise basal kinase activity two-fold (Busch et al., 2002), presumably due to loss of the S80 hydroxyl contacts with Q417 and A81 and the associated destabilization of the R:C interface.

The interdomain helices and the CNB-B αA helix shield the rest of the activation loop from solvent though hydrophobic and hydrogen bonding interactions (left zoom-in panel of site 3 in ***Figure 2B***). The extended interdomain helices dock to one side of the activation loop while the CNB-B αA helix docks to the curved tip of the activation loop (right zoom-in panel of Site 3 in ***Figure 2B***). The side chains of M216, M217 and H224 at one side of the interdomain helices make van der Waals contacts with side chains of K529 and W531 at the activation loop. The side chain of E243 in the CNB-B αA helix is oriented parallel to the curved tip of the activation loop and docks to F526 and G527 through van der Waals contacts while the side chain of D250 forms a salt bridge with the side chain of K529 (right zoom-in panel of Site 3 in ***Figure 2B***). This region differs substantially from previously reported PKG structures: in the cGMP-bound structure of the R-domain (Kim et al., 2016), the interdomain linker consists of bent helices that make R:R intermonomer contacts.

At site 4, the loop between the αH and αI helices at the bottom of the C-domain interacts with the αB helix of CNB-B in the R-domain exclusively through hydrophobic contacts (***Figure 2A***) to bury a small hydrophobic surface area of 103 Å^2^. As seen in the bottom panel of ***Figure 2B***, a hydrophobic cluster consisting of H338, L339 and I340 at the end of the αB helix docks to S615, L621 and L622 in an S-shaped loop (residues 615-624). Because the CNB-B αB helix is tightly coupled to the PBC, conformational changes that occur upon cGMP binding can disrupt the above interactions at site 4 and enable activation.

### C-domain architecture

The PKG Iβ C-domain in the auto-inhibited complex is very similar to a structure of the isolated C-domain (Qin et al., 2018), showing a RMSD of 0.39 Å for 322 Cα atoms that exclude the glycine rich loop, which is displaced by up to 3.5 Å in the isolated C-domain to accommodate a bound inhibitor (***Figure supplement 4***). While the overall fold of the auto-inhibited PKG C-domain is similar to that of the auto-inhibited PKA catalytic subunit, differences in the orientations and contacts of the αA helix and the C-terminal tail of PKG compared to PKA may be important to understanding how signaling pathway cross-talk is minimized and how the mechanisms of activation differ (***Figure 3A***). In PKG, the αA helix interacts only with the small lobe, whereas the αA helix interacts mainly with the large lobe in the C-subunit of PKA. As shown in ***Figure 3B***, the PKG αA helix is linked to the adenosine pocket via the E363:R438 salt bridge and F367:Y440 packing interaction. The significance of this difference is unclear, since residues that precede the αA helix in the auto-inhibited complex are not ordered and show no density (residues 330-353).

**Figure 3.**
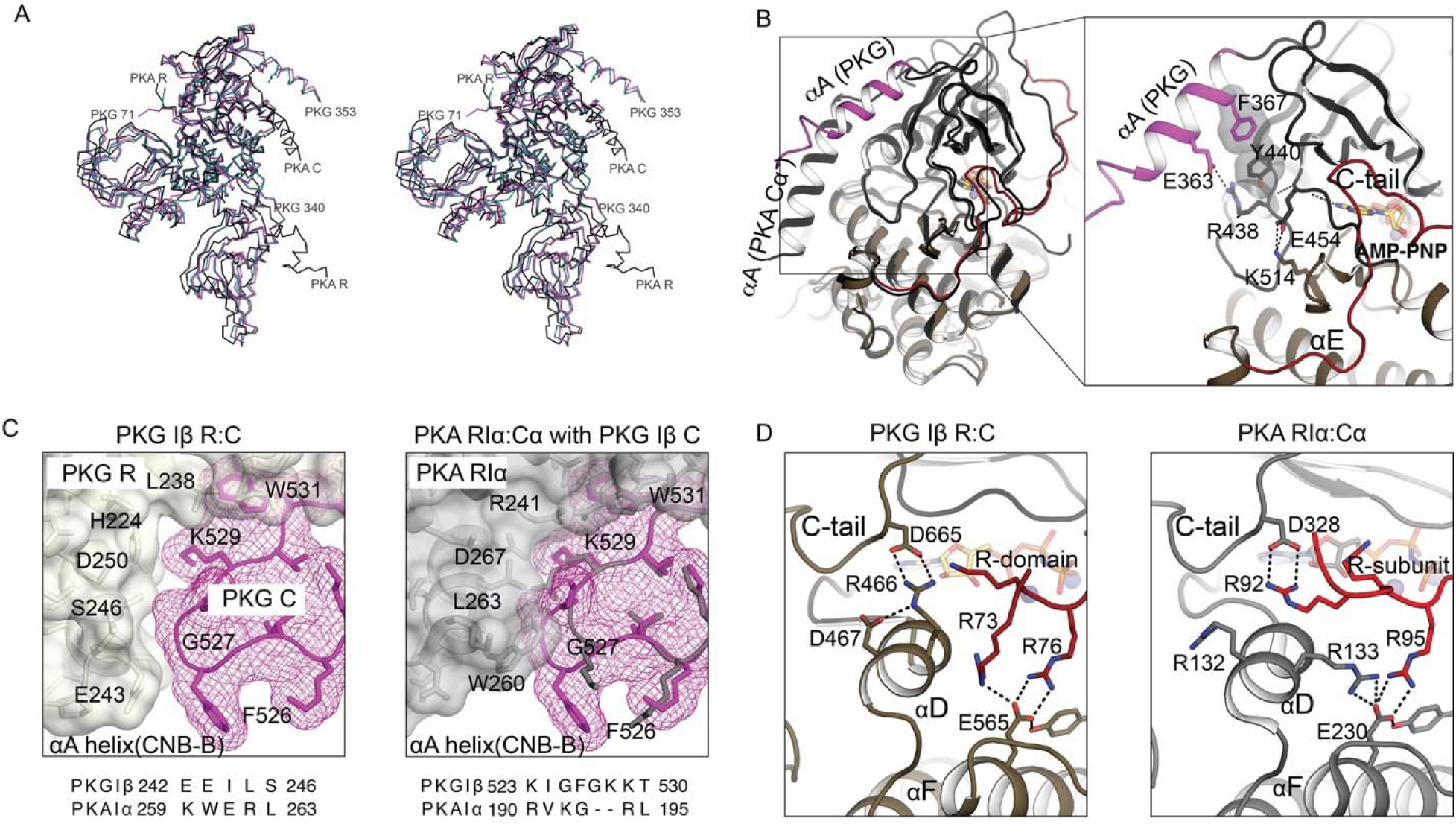
Comparison of auto-inhibited PKG and PKA structures. (A) Structural alignment of auto-inhibited PKG Iβ (red, PDB ID: 7LV3) and PKA Iα (gray, PDB ID: 2QCS) in stereo-view. (B) C-domain αA helices are differently positioned in PKG Iβ and PKA. Left: auto-inhibited PKG Iβ and PKA Iα structures are aligned using Cα atoms of the C-domains Right: Zoom in view of interactions between the PKG Iβ αA helix and the catalytic core. The interacting residues and bound AMP-PNP:Mn^2+^ (ANP) are shown in sticks and labeled, and hydrogen bonds are shown as dotted lines. The short PKG Iβ αA-β1 loop allows αA to contact the small lobe of the C-domain via a salt bridge (E363:R438) and stacking interactions (F367:Y440) in a way that may be conveyed to the ribose pocket. (C) PKG Iβ contacts between the CNB-B αA helix and the C-domain near the activation loop imply that binding of PKA RIα would cause steric clashes. Left: PKG Iβ R:C interface at Site 3 with sequences of PKG Iβ and PKA Iα at the CNB-B αA helix aligned. The activation loop is shown in mesh (magenta), the R-domain is shown in transparent surface (grey), and interacting residues are labeled. Right: Aligning auto-inhibited PKA Iα on auto-inhibited PKG Iβ using the C-domains (except the activation loop) shows that large PKA RIα side chains L263 and W260 that contact the shorter PKA Cα loop would clash with the larger PKG Iβ loop. Sequence alignment between PKG Iβ and PKA Iα at the activation loop is shown below. (D) In auto-inhibition, PKG Iβ and PKA C-terminal tails occupy similar positions but contact different partners. The C- and R-domains (or subunits) are colored in grey and red, respectively. Left: The C-tail of PKG interacts with the C-domain αD helix and the AI linker interacts with the C-domain αF helix. Right: The C-tail of PKA Iα interacts with the AI linker, and both the αD helix and the AI linker interact with the C-subunit αF helix.

Compared to PKA C, PKG Iβ has a longer activation loop connecting αA and β1 due to a two amino acid insert. Superimposing the PKG and PKA structures (excluding loop residues) shows that the extended PKG activation loop is accommodated by small PKG R-domain residues E243 and S246, whereas the PKA activation loop contacts larger PKA RIα residues W260 and L263 at the homologous positions. We infer that PKA RIα binding to the PKG C-domain would cause clashes: the extended tip of the activation loop of PKG I would clash with the CNB-B A-helix of PKA RIα (***Figure 3C***). These sequence differences, and those of the AIs (***Figure 1C***) discussed above, likely arose from evolutionary pressure to minimize cross-talk between the cyclic nucleotide signaling pathways.

The C-terminal tails of AGC kinases contribute to the overall fold and catalytic function (Kannan et al., 2007) and is important for kinase assembly and regulation (Taylor et al., 2021). In PKG, the C-terminal tail interacts directly with the kinase core (***Figure 3D***): D665 of the C-tail forms a strong salt bridge with R466 of the αD helix, and D467 stabilizes the position of R466 with a hydrogen bond. The residue at the corresponding position in the PKA C-terminal tail is also an aspartate (D328), but it interacts with R92 of the RIα subunit (P-5), rather than with an arginine in the αD helix of the C-subunit. In PKA, R133 of the αD helix (which is homologous to R466 of PKG Iβ) points towards the active site and interacts with E230 of the αF helix, perhaps because the PKA C-subunit has a preceding arginine (R132) rather than an aspartate (D467 in PKG Iβ C-domain) that causes charge repulsion.

### Conformational differences between auto-inhibited and activated states

The conformation of the PKG Iβ AI has not previously been described, but hydrogen/ deuterium exchange experiments have shown that amide backbone hydrogens in the AI domains of both PKG Iα and PKG Iβ exchange rapidly in the presence of cGMP, indicating that this region is highly dynamic (Alverdi et al., 2008; Chan et al., 2020), so we expect that the AI is likely to be disordered when not bound to the catalytic cleft. The contacts made by the AI in this structure may contribute to the allosteric activation mechanism by recruiting residues of PBC-A and the αB helix that form the docking surface for the C-domain.

Comparing the auto-inhibited complex with previous structures of PKG R-domain fragments bound to cyclic nucleotides (Kim et al., 2016; Osborne et al., 2011) reveals local changes in each CNB domain, as well as global changes that orient the two CNBs so that they clamp around the large lobe of the C-domain (***Figure 4A***). These R-domain changes map primarily to the helical subdomains, especially the interdomain helices, which become extended and position CNB-A and -B relative to each other for binding to the C-domain (***Figure 4B***). We also observe conformational differences between the activated and inhibited states within the CNB domains themselves. Previous studies in other systems, including PKA, have shown that CNB β-subdomain structures are largely invariant but that ligand binding can induce changes in the PBC, which adopts an ‘open’ apo state but a ‘closed’ state with bound nucleotide (Berman et al., 2005; Kim and Sharma, 2021; Taylor et al., 2008). Tracking the PBCs (and adjacent helices) enables us to develop a rationale for how structural changes induced in the CNBs are coupled to the stabilization of the PKG auto-inhibited state.

**Figure 4.**
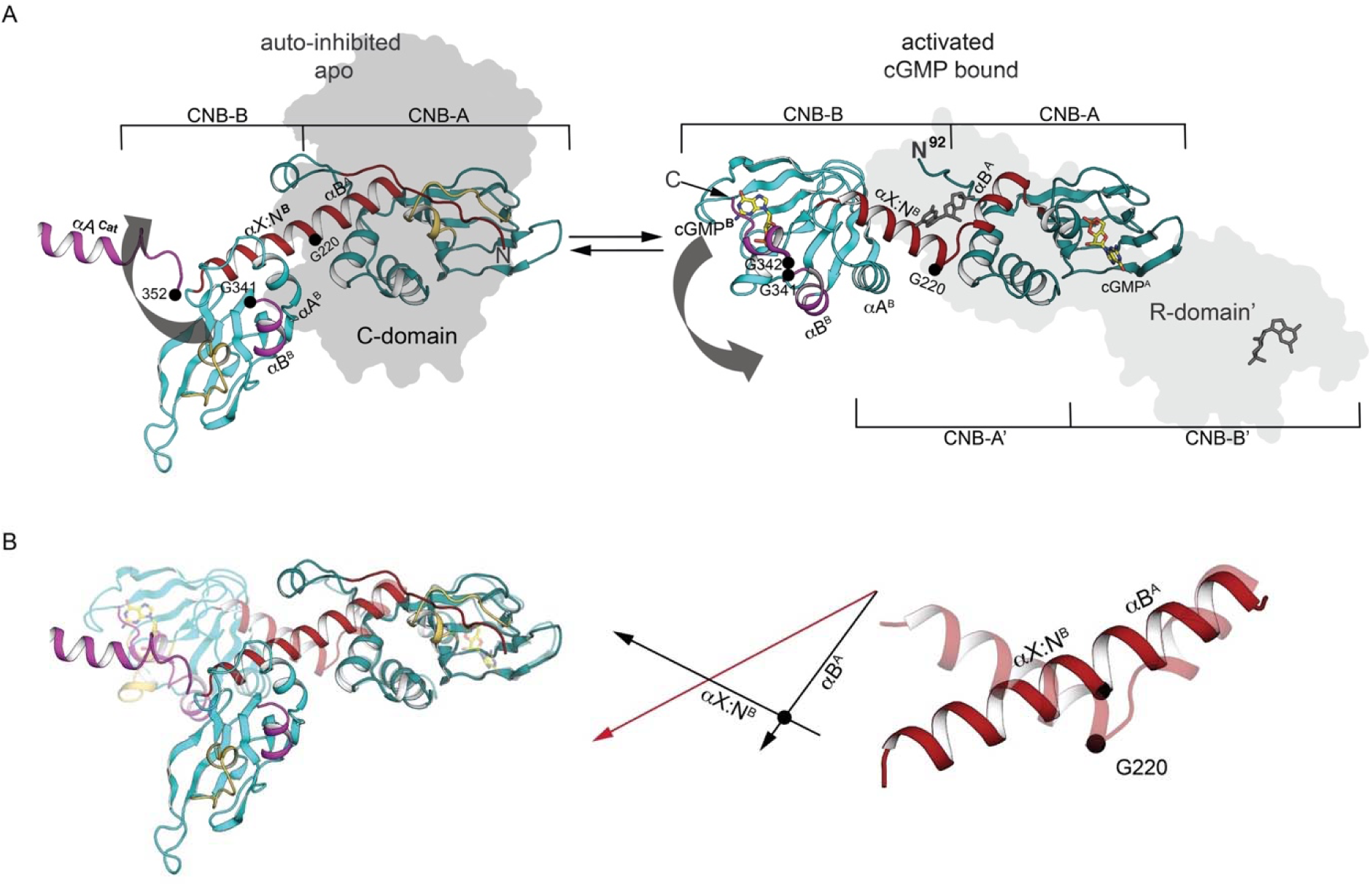
Comparison of PKG Iβ R-domain structures in activated and auto-inhibited states. (A) Cartoon of a single R-domain in either (left) the auto-inhibited R:C state, bound to the C-domain depicted as a grey silhouette, or (right) the activated R:R state, bound to cGMP and another R-domain shown as a light grey silhouette (PDB ID: 4Z07). The interdomain helices are labeled and shown in red, and G220, G341, and G342 are marked. (B) Structural alignment of interdomain helices in the auto-inhibited and activated conformation. Left: The R-domain in the auto-inhibited state (solid) and activated state (transparent) conformations are aligned using CNB-A (at right). Middle: orientations of the interdomain helical axes in the auto-inhibited state (red) and the activated state (black). Right: cartoon depiction of the interdomain helices only, with a blck circle indicating the position of the G220 Cα atom in both the auto-inhibited state (solid) and the activated state (transparent).

The β-subdomain of CNB-B in the auto-inhibited holoenzyme superimposes well on the structure of the cGMP-bound isolated CNB-B domain, but the CNB-B PBC adopts an ‘open’ state in the former and a ‘closed’ state in the latter (***Figure 5A***). With respect to previously determined structures, CNB-B in the auto-inhibited holoenzyme is most similar to the structure of the isolated apo CNB-B domain structure (RMSD of 0.88 Å for 106 shared Cα atoms, ***Figure supplement 5***); the auto-inhibited domain is somewhat more open than the isolated apo domain. The CNB-B αC helix ends at the disordered 15 residues for which the crystal has no density, so we do not interpret its contacts here.

**Figure 5.**
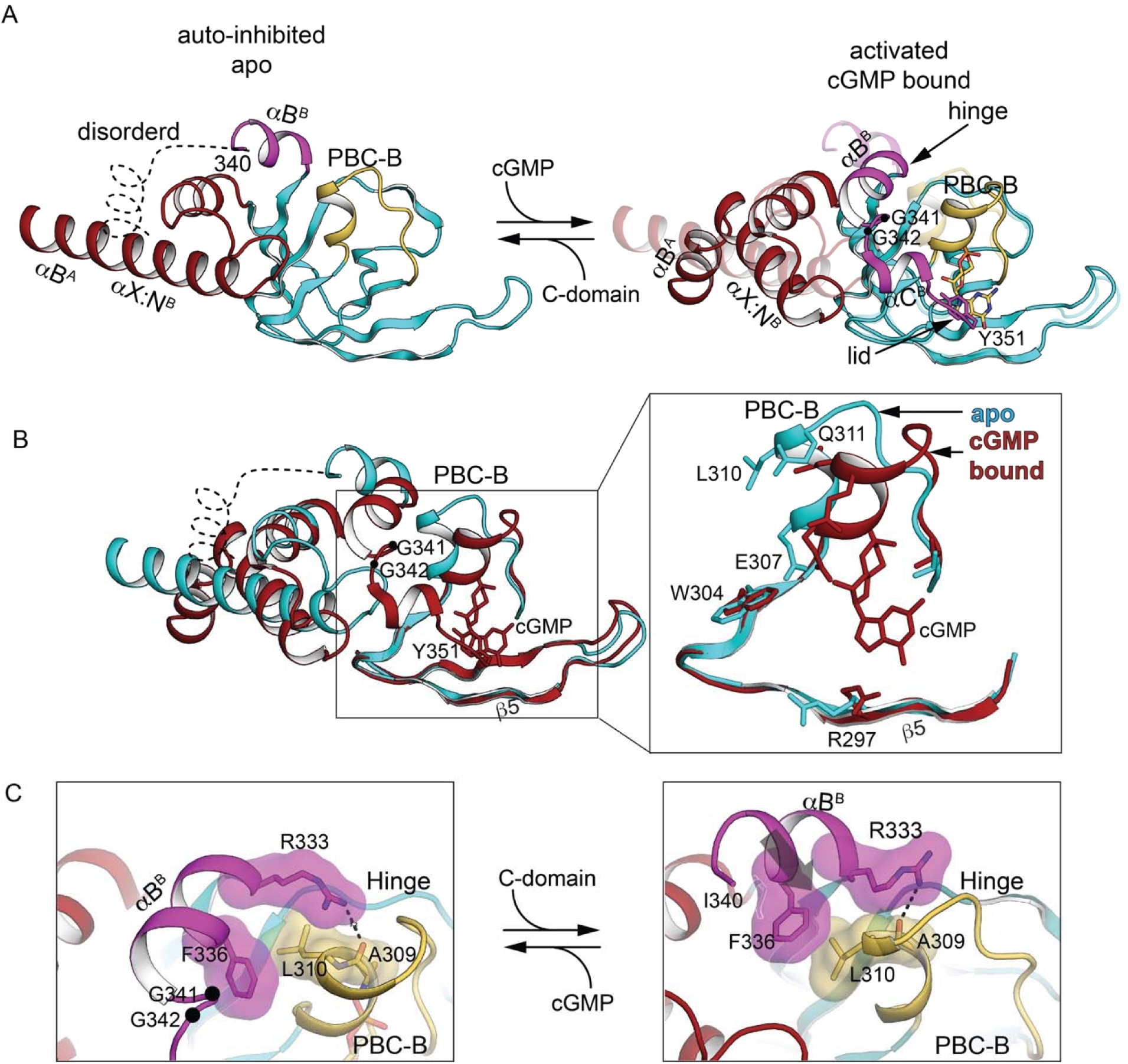
The CNB-B helical subdomains are structurally distinct in auto-inhibited and activated conformation. (A) Overall structure of the interdomain helices and CNB-B in the auto-inhibited (left, PDB ID: 7LV3) and superposition of the cGMP-bound, activated state of the tandem CNB dimer (solid) and auto-inhibited (transparent) (right, PDB ID: 7LV3 and 4Z07). Hinge glycine Cα atoms are labeled and cGMPs are shown in stick. In the cGMP-bound ‘closed’ state, PBC-B moves along with αB^B^ helix and the hydrogen bond interactions between αC helix (lid) and PBC-B provide cGMP capping interactions. (B) Superposition of CNB-B in the tandem CNB cGMP-bound dimer (red, PDB ID: 4Z07) and in the auto-inhibited complex (cyan, PDB ID: 7LV3) using the invariant β domain elements. Left: β4, β5, and the β4-β5 loop superimpose well (bottom) but the interdomain helices adopt very different conformations. Right: Zoomed-in view of the differences between the PBC loops of the two states. In the auto-inhibited ‘open’ state, the αB helix of CNB-B docks to the C-domain, the αC helix is disordered, and the more open PBC-B interacts directly with N3A. (C) Zoomed-in view of the hinge region. Key hinge residues are shown in stick with transparent surface.

The differences between CNB-B in the auto-inhibited complex described here and the previously reported cGMP-bound R:R dimer (Kim et al., 2016) resemble the differences between the inhibited and activated states of CNB-A of PKA RI (Kim et al., 2005). The β-sandwich is unchanged while the PKG Iβ PBC is ‘open’ in the apo, auto-inhibited state and ‘closed’ in the activated states (***Figure 5A***). In the auto-inhibited state, the αB^B^ helix docks to the C-domain (***Figure 2B***, Site 4) in a very different orientation relative to the β-subdomain than is seen in the cGMP-loaded state (***Figure 5B***). N3A^B^ contacts the open PBC^B^, filling the space that the αC^B^ helix occupies in the cGMP-bound state. The αC^B^ helix, the C-loop, and the capping residue Y351 are disordered, whereas in the activated state the hinge formed by residues G341 and G342 of the B helix positions αC^B^ against PBC^B^ and Y351 shields cGMP from solvent (***Figure 5A***). Although the orientation of PBC^B^ with respect to the β subdomain differs substantially between the auto-inhibited and cGMP-bound states (***Figure 5B***), A309 and L310 at the tip of PBC^B^ interact with R333 and F336 of the αB^B^ helix in both states, acting as a hinge.

CNB-A in the auto-inhibited PKG Iβ holoenzyme adopts an ‘open’ conformation typical of apo CNBs (***Figure 6A and B***), and its contacts with the C-domain indicate that the orientation of the PBC is critical to auto-inhibition. Superimposing a structure of the cGMP-bound isolated CNB-A domain with the R:C complex CNB-A shows that L184 and Y188 in the PBC-A helix would clash with M579 and Y583 of the C-domain αG helix, so the ‘closed’ state is sterically incompatible with binding to the C-domain (***Figure 6C***). In the context of an R-domain dimer, however, the ‘closed’ PBC of cGMP-bound CNB-A contacts the αB helix of CNB-A in another monomer (***Figure 6D***) (Kim et al., 2016). Thus, cGMP binding stabilizes a CNB-A PBC conformation that cannot support R:C interactions but does support R:R interactions. Other CNB-A residues at the R:C interface make alternate interactions in the activated R:R dimer (***Figure 6D***). At the R:C interface of the auto-inhibited state, Y188 (PBC-A) and Q213 (αB-helix) interact with I79 and P83 of the AI, and M217 (αB-helix) docks to the activation loop residue W531 (right panel in ***Figure 6C***), but in the cGMP-bound activated R:R dimer, Y188 and Q213 from one chain interact with M217 and N189 from the other chain (left panel in ***Figure 6D***).

**Figure 6.**
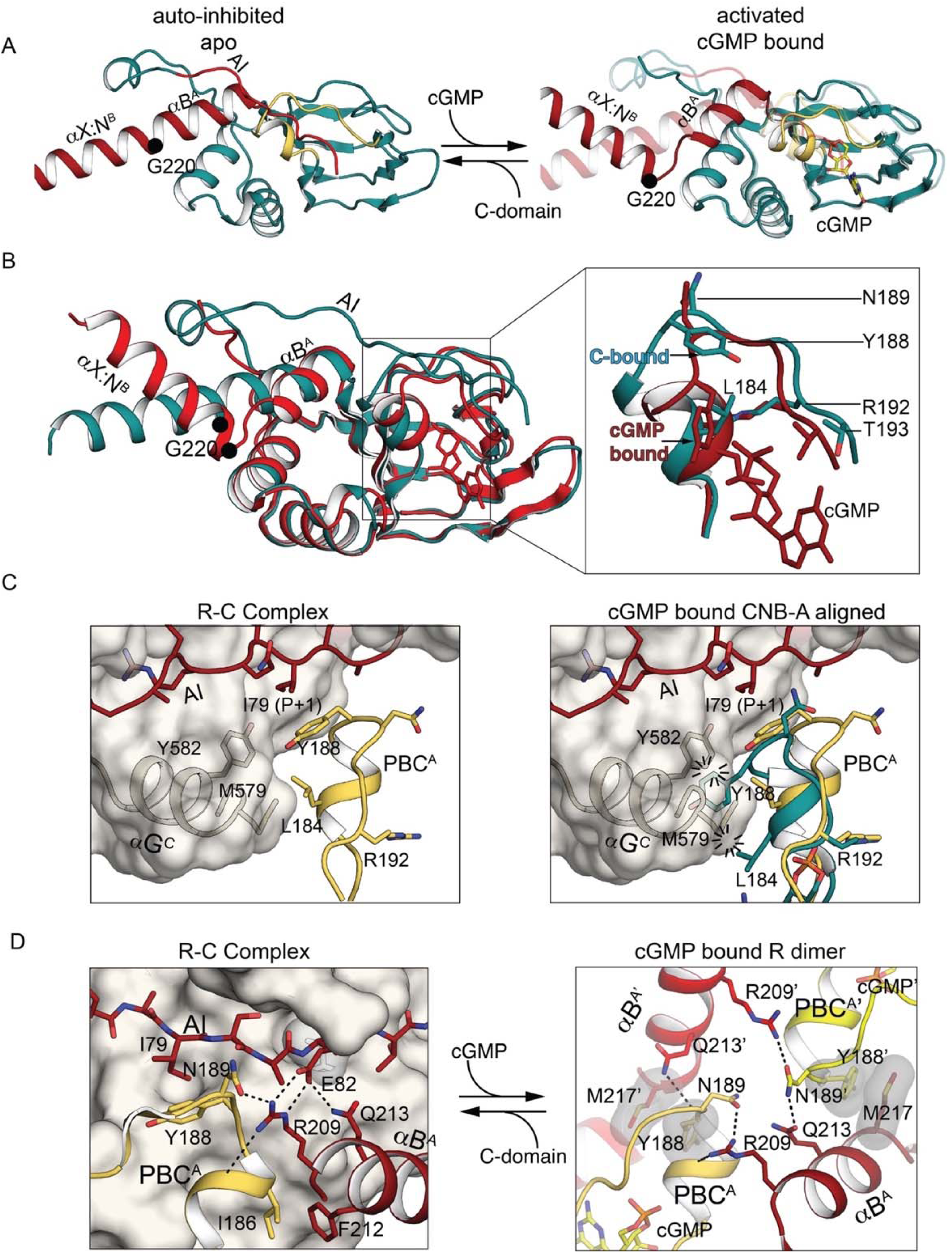
CNB-A PBC and inter-domain linker differences in auto-inhibition compared to activation. (A) Overall structure of the interdomain helices and CNB-A in the auto-inhibited (left, PDB ID: 7LV3) and superposition of the cGMP-bound, activated state of the tandem CNB dimer (solid) and auto-inhibited (transparent) (right, PDB ID: 7LV3 and 4Z07). The same color scheme is used as in Figure 2. The Cα atom of G220 is marked with a sphere. (B) Structural alignment of cGMP-bound (PDB ID: 4Z07) and C-domain-bound conformation (PDB ID: 7LV3). CNB-A in cGMP-bound and C-bound conformations are colored in red and dark teal, respectively. Zoomed-in view at the right panel shows PBC and αB helix. Key hinge residues and R:C interface residues are shown in stick. (C) Alignment of CNB-A:cGMP with PKG holo complex. Left: R:C interface near the apo PBC-A is shown. PBC-A and the αG helix are shown in cartoon. The C-domain is shown as transparent surface. Key R:C interface residues are shown in sticks. Right: Structural alignment between CNB-A:cGMP with PKG Iβ holoenzyme. The residues L184 and Y188 of PBC-A of the cGMP-bound state show steric clashes with M579 and Y583 of the G helix at the C-domain and suggests that CNB-A:cGMP is not compatible with the C-domain. (D) Dynamic AI region replaces the R-R contacts upon the R:C complex formation. Left: The R-R dimer interface near PBC-A is shown. Residues from the second chain are marked with ‘. Right panel shows AI, PBC-B and B helix of the R-domain in cartoon and the C-domain in surface. R209 helps assemble a docking surface consisting of AI, PBC-B and B helix of the R-domain.

Located at the C-terminal end of the CNB-A β subdomain, the αB helix functions as a flexible hinge that allows conformational changes required for cyclic nucleotide binding and release through a conserved Phe or Tyr that contacts the PBC through hydrophobic interactions (Berman et al., 2005; Rehmann et al., 2007). In the auto-inhibited state, I186 of the open PBC lies in a hydrophobic core formed by F212 of the CNB-A αB helix and F118, M119, and V155 (left zoom-in panel of site 2 in ***Figure 2B***), but in the activated state I186 is displaced from this core and these contacts are lost in the closed PBC. The PBC undergoes a substantial but very localized structural change between these two states that reorganizes the tip of the PBC: the Cα atoms of residues 185 through 189 are displaced by 3.4 to 7.5 Å (5.5 Å RMSD), and the Y188 side chain is displaced by up to 8.3 Å, but flanking residues such as R192 are only minimally displaced (right zoom-in panel in ***Figure 6B***) (Berman et al., 2005; Kim and Sharma, 2021; Taylor et al., 2008).

### Structural basis for a PKG I activating mutation

Mutation R177Q in PKG Iα is associated with TAAD and constitutively activates the holoenzyme despite weakening cGMP affinity for the isolated CNB-A domain more than 10^5^-fold (Guo et al., 2013). Introducing R192Q into PKG Iβ is also activating (Chan et al., 2020). In the auto-inhibited state of PKG Iβ, the R192 guanidinium group donates hydrogen bonds to the L154 and G182 carbonyl oxygens; in the isolated cGMP-bound CNB-B domain (Kim et al., 2011), it hydrogen bonds to these oxygens and also to a non-bridging oxygen of the cyclic phosphate in the cGMP-bound domain (***Figure supplement 6A***). Loss of this key cGMP contact explains the decreased cGMP affinity of the isolated RQ domain, but it is not obvious why the intact enzyme should be activated. Since CNBs canonically adopt ‘open’ conformations in the apo state and ‘closed’ conformations with bound cyclic nucleotide, loss of the conserved arginine could cause activation if its contacts to the L154 and G182 carbonyl oxygens stabilize the open state more than the closed state.

We determined the structure of the RQ variant of the isolated CNB-A domain (PDB;7MBJ), revealing that it adopts a closed conformation and closely mimics that of the cGMP-bound wildtype isolated domain (***Figure supplement 6B and 6C***). Since the RQ mutation does not alter the folding of the domain or prevent it from adopting the closed conformation, we propose that the rest of the sequence of PKG I CNB-A is biased to adopt the closed state. Such bias would explain why CNB-A binds cGMP 18-fold tighter than CNB-B despite largely conserved contacts (Kim and Sharma, 2021): CNB-B samples the open state more strongly, so more free energy of binding is needed to drive the equilibrium to the closed state. Skewing of the equilibrium between open and closed states has also been proposed to explain the 3-order of magnitude difference in cAMP affinities for PKA RIα and the CNB domain of HCN2 (Moleschi et al., 2015).

Because the PBC of the RQ CNB-A domain makes intermonomer crystal contacts, it is possible that these contacts help stabilize the closed state. To determine if the closed state would persist in the absence of crystal contacts, we performed molecular dynamics (MD) simulations of the fully solvated domain without nucleotide, and of the wildtype domain with or without nucleotide (***Table supplement 3***). Analyzing the coordinates from these MD trajectories shows that all three domains sample conformational space that is more similar to the closed state (cGMP-bound wildtype CNB-A, PDB 3OD0) than to the open state (CNB-A in the auto-inhibited structure) (***Figure supplement 7A***) (VanSchouwen et al., 2015a). We conclude that neither crystal contacts nor interactions with cGMP are needed to drive either the wildtype or mutant domain to the closed state, supporting our hypothesis that the CNB-A domain is biased toward the closed state.

We previously showed that binding of cyclic nucleotide induces backbone amide NMR chemical shift perturbations in CNB domains that reflect the degree to which the closed state is populated (Akimoto et al., 2015; Byun et al., 2020). To experimentally assess the degree to which the isolated R192Q CNB-A domain might sample the closed state in solution, we acquired NMR spectra of the isolated wildtype domain in the apo and cGMP-loaded states, and of the apo R192Q mutant (***Figure supplement 8***). Chemical shifts of apo R192Q mutant NH resonances are more similar to those of the cGMP-bound wildtype domain than to the apo wildtype domain (***Figure supplement 8***), consistent with the mutation inducing a conformational change similar to that induced in the wildtype domain by cGMP binding. We therefore conclude that the RQ mutation causes constitutive kinase activation by allowing the CNB-A domain to adopt a closed conformation similar to that of cGMP-bound wildtype without binding cyclic nucleotide.

### PKG Iβ is auto-inhibited in cis

While the structure in the crystal likely represents a domain-swapped dimer (Gronenborn, 2009) undergoing auto-inhibition in trans, previous biochemical studies of truncated PKG lacking the LZ dimerization domain have all been consistent with inhibition in cis. PKG Iβ lacking the LZ domain elutes from gel filtration columns as a monomer (Richie-Jannetta et al., 2003), and we found that a monomeric PKG Iβ lacking the first 55 amino acids has low basal activity in the absence of cGMP, showing that the monomer is capable of auto-inhibition (Kim et al., 2016). Because it is formally possible that auto-inhibition of the native dimeric enzyme might occur in cis or in trans, we used engineered full-length PKG Iβ heterodimers to assess any roles of interchain communication in regulating PKG activity (Chan et al., 2020). In this method, cells are co-transfected with an untagged ‘active’ PKG and with a Flag-tagged ‘dead’ PKG bearing a point mutation that renders the kinase domain catalytically inactive. When protein is isolated with anti-Flag agarose and eluted with Flag peptide, the resulting purified kinase is a mixture of dead-dead homodimers and dead-active heterodimers; in assays, activity comes only from the heterodimers. In this system, introducing mutations in the dead chain cannot affect the kinase activity of that chain (intrachain effect), since the active site is mutated, so any changes in activity must result from interchain effects. We prepared constructs that mutated K75 and R76 to glutamate (KR/EE), reasoning that these AI residues make key R:C contacts with acidic residues at the active site, so this change would cause charge-repulsion and destabilize the auto-inhibited R:C state.

We performed kinase assays under basal conditions (no cGMP) and activating conditions (10 μM cGMP) using Flag-tag purified protein from four separate co-transfections that pair Flag-tagged kinase-dead PKG Iβ (with or without the KR/EE mutation) with untagged kinase-active PKG Iβ (with or without the KR/EE mutation) (***Figure 7***). With wild-type active chain, low levels of activity are seen in the absence of cGMP, indicating auto-inhibition, and high levels of activity are seen with cGMP, consistent with release of auto-inhibition. When the KR/EE mutation is present on the active chain, similar high activity levels are seen with or without cGMP, indicating that little or no auto-inhibition occurs. We infer that native dimeric PKG Iβ undergoes auto-inhibition primarily in cis. We note that the activity of the KR/EE mutant active chain is slightly lower when paired with a wildtype Flag-tagged partner than when paired with the AI mutant, indicating that the wild-type AI may provide some modest inhibition in trans. The basal kinase activity of the untagged wild-type chain is slightly higher when it is associated with the dead chain bearing the KR/EE mutation (5.4% of maximum) rather than the wild-type AI (3.1% of maximum), suggesting that the presence of a second copy of the AI in trans may modestly enhance auto-inhibition of the engineered heterodimer.

**Figure 7.**
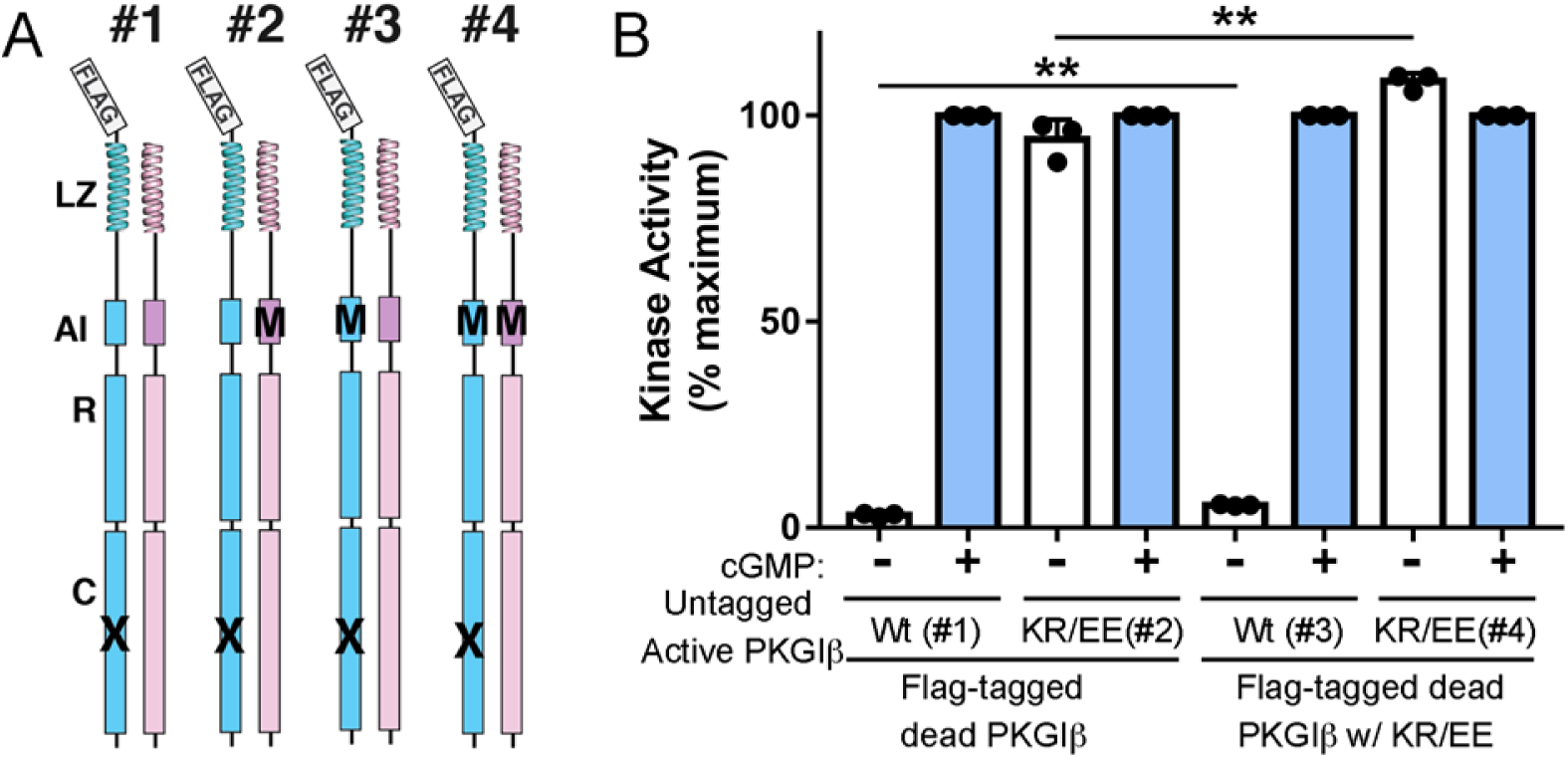
The PKG Iβ AI sequence does not contribute to activation through interchain contributions. (A) Anti-Flag pulldown of a Flag-tagged (blue) catalytically ‘dead’ PKG Iβ (X) permits isolation of heterodimers containing an untagged ‘active’ PKG Iβ (pink) from co-transfected cells. Flag-tagged homodimers that are also present have no kinase activity. Introducing AI mutations K75E and R76E (KR/EE in the text, shown schematically as M) in the ‘dead’ and/or ‘active’ chain allows us to distinguish intrachain and interchain influences on auto-inhibition. (B) Kinase activity for four heterodimers at zero cGMP (-) or 10 μM cGMP (+) show that AI mutations in the active monomer (heterodimers 2 and 4) confer constitutive activity and eliminate auto-inhibition. Presence of the AI mutation in the ‘dead’ monomer alone (heterodimer3) does not eliminate auto-inhibition, though it may slightly enhance basal activity.

### PKG Iβ activation is potentiated by substrate

Our structure indicates that mutations such as S80A (Busch et al., 2002) or KR/EE promote cGMP-mediated activation by weakening interactions between the AI sequence and the active site cleft, and thus destabilizing the auto-inhibited R:C state. Interestingly, the PKG Iβ cGMP activation constant can be modulated by the peptide substrate: PKG Iβ activates at two-fold lower cGMP levels with VASPtide as a substrate than with Kemptide (Figs. 1D and S3). This observation is consistent with a competition between the AI pseudo-substrate and the actual substrate for accessing the catalytic cleft, with binding of substrate preventing formation of the auto-inhibited state; the two-fold lower *K*_M_ for VASPtide compared to Kemptide (***Figure supplement 3***) explains enhanced activation by the former over the latter. Competition of substrate with the AI sequence for binding to the catalytic domain has also been invoked in explaining the activation of PKA (Byun et al., 2020).

In comparison with PKG Iβ, PKG Iα exhibits higher fractional basal activity, a lower activation constant, and only small differences in cGMP activation constants with Kemptide and VASPtide (***Figures 1D and Figure supplement 3***). Such differences may reflect not only poorer contacts between the Iα AI sequence and the conserved active site (as discussed above for Site 1 of ***Figure 2B***), but also more limited access of the Iα AI to the kinase active site cleft compared to the Iβ enzyme. The Iα-specific leucine zipper, linker, and hinge regions that contribute to cGMP-mediated activation (Ruth et al., 1997) might combine to sterically restrict (relative to PKG Iβ) how AI diffusion samples its environment to access the kinase active site cleft. The construct crystallized here lacks the leucine zipper and linker, so we have no detailed basis for explaining how such restriction might arise; structures of full-length proteins may be needed to clarify these isoform-specific aspects of the activation mechanism.

## Discussion

### PBC conformations control access to the auto-inhibited state

Based on structures of the isolated CNB-B domain, we previously proposed that cGMP drives release of auto-inhibition by forming hydrogen bonds with the cGMP phosphate that induce the PBC loop to close and reposition the adjacent helix (***Figure supplement 7***) (Campbell et al., 2017; Huang et al., 2014a; Huang et al., 2014b). In the auto-inhibited PKG Iβ 71-686 structure presented here, the PBC loop of CNB-B and its adjacent helix adopt conformations very similar to those of the isolated apo domain; this arrangement allows αA- and αB-helices to contact the C-domain, whereas the repositioning of these helices as in the cGMP-bound closed state would disrupt these contacts (***Figures 2B and 5***), providing further support for our hypothesis. NMR data show that the isolated apo CNB-B domain is predominantly open but samples the closed state (VanSchouwen et al., 2015b). The closed state is stabilized by cGMP binding, while the PKG Iβ 71-686 structure provides a snapshot of the contacts that stabilize the open state of apo CNB-B and support auto-inhibition. CNB-B discriminates strongly between cGMP and cAMP (Huang et al., 2014b), and certain contacts are critical to confer the conformational change, as highlighted by the structure of the isolated CNB-B domain in an inhibitor-bound open state (Campbell et al., 2017; Huang et al., 2014a; Huang et al., 2014b): although *R*_P_-cGMPS makes many of the same contacts as cGMP, the PBC loop does not close to contact the phosphorothioate sulfur, and the adjacent helix does not reposition (***Figure supplement 9***). Whereas cGMP binding to PKG Iβ CNB-B disrupts its αA- and αB-helix contacts with the C-domain, resulting in activation, we expect that *R*_P_-cGMPS binding will not disrupt the R:C interface of the auto-inhibited state, explaining why this cGMP analog acts as an inhibitor.

The open state of CNB-A also supports favorable R:C contacts in auto-inhibited PKG Iβ that would be incompatible with the closed state that is stabilized by binding cGMP (***Figure 4***). The auto-inhibited structure presented here is the first instance in which a fully open state of PKG I CNB-A has been observed; if the isolated CNB-A domain samples the open and closed states, as CNB-B does (Huang et al., 2014b), the tighter cGMP affinity of CNB-A compared to CNB-B (Kim et al., 2011) could be explained by apo CNB-A being biased strongly to the closed state whereas CNB-B is biased to the open state. Similar coupling of cyclic nucleotide binding to differently skewed equilibria between the open and closed states has been proposed to explain differences in cAMP affinities for PKA RIα and the CNB domain of HCN2 (Moleschi et al., 2015). We propose that the open CNB-A domain seen in the auto-inhibited enzyme is a high energy state stabilized by R:C contacts that rapidly reverts to the closed state when these contacts are lost. The auto-inhibited structure provides a basis for understanding how two analogous CNB domain switches with very different intrinsic setpoints are coupled to one another in an allosteric mechanism that exhibits one macroscopic activation constant. A quantitative description of this coupling may require investigating the local and global dynamics of the enzyme.

### Activation mechanism

We propose a model for PKG Iβ activation that combines available biochemical data to posit roles for several distinct enzyme substrates that we relate to known crystal structures (***Figure 8***). At low cGMP concentration, the AI sequence and the helical sub-domains of the R-domain dock to the large lobe of the C-domain as in our auto-inhibited complex, blocking the active site and completely shutting off kinase activity (***Figure 8***, bottom left). Phosphorylated threonine 532 of the C-domain activation loop stabilizes the docking surface for the R-domain by forming a hydrogen bonding network involving both C- and R-domain residues as seen in ***Figure 2B***.

**Figure 8.**
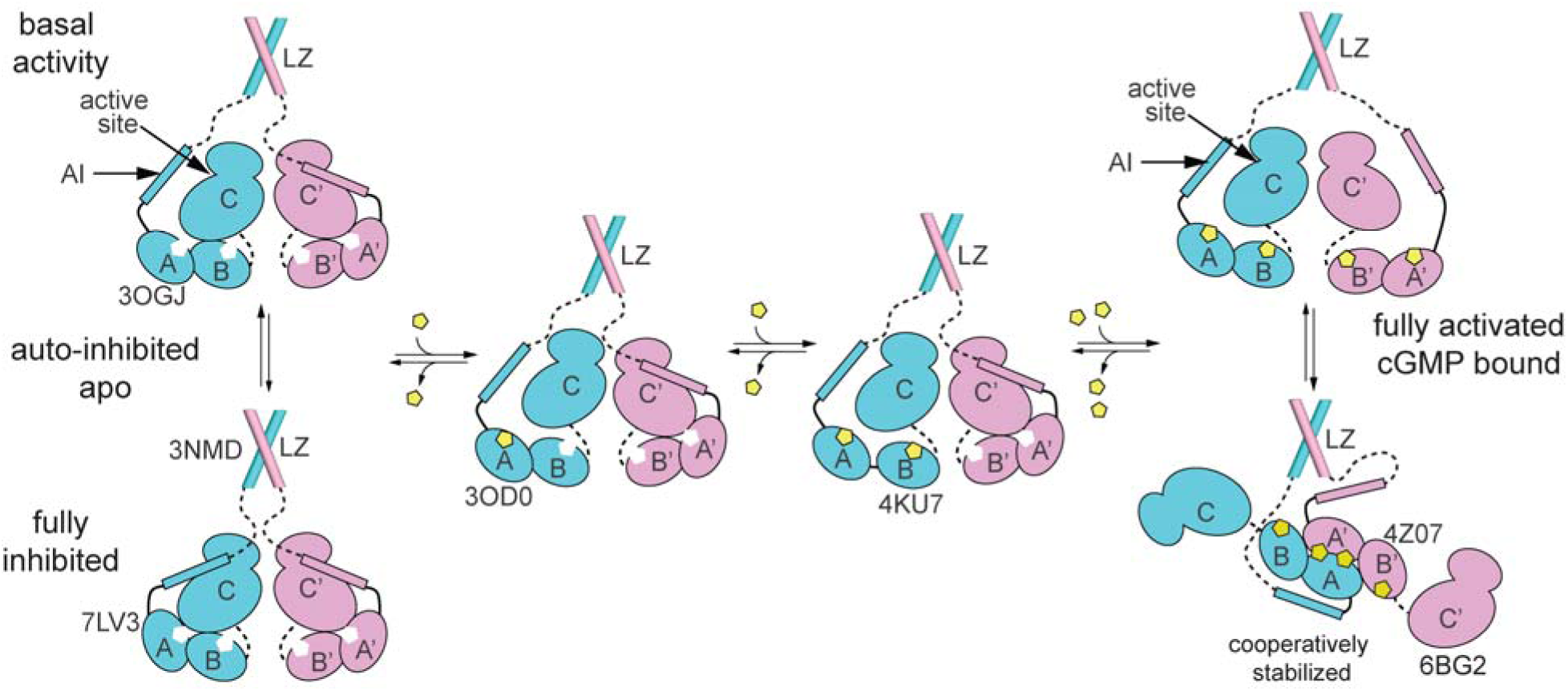
Schematic model for how equilibria between different states of PKG Iβ support auto-inhibition and activation. Crystal structures that represent one or more domains in each state are indicated by PDB code. The auto-inhibited state (bottom left) dimerizes through leucine zipper domains (3NMD) that keep the cGMP-free, auto-inhibited catalytic domains (7LV3) in close proximity. The AI sequence of cGMP-free PKG Iβ can transiently leave the active site and permit basal activity (top left). Binding of cGMP to CNB-A (middle left; 3OD0) and/or CNB-B (middle right, 4KU7) lock the CNBs into conformations that are not compatible with the auto-inhibited state. Levels of cGMP that saturate CNBs of both monomers enable complete activation (top right) and support formation of a complex between the tandem CNB domains that provides cooperativity and may restrict the ability of the AI sequence to access the catalytic active site. The discussion further describes how peptide substrate and cGMP analogs, including cAMP, can influence the equilibria.

Basal kinase activity in the absence of cGMP occurs during transient excursions of the AI away from the active site cleft (***Figure 8***, top left). These excursions explain the slow but complete hydrogen/deuterium exchange of AI backbone amides in the wild type apo PKG Iβ (Chan et al., 2020). The frequency of these excursions should depend on the AI sequence and phosphorylation state, which influence the detailed interactions between the AI and the C-domain as discussed above. Phosphorylation of S80 during an excursion would strongly bias against a return to the auto-inhibited state, and occupation of the active site by a competitor would prolong the excursion. Since PKG Iβ shows about 40-fold activation (***Figure 1D and Figure supplement 3***), in the absence of cGMP we expect that transient excursions expose the kinase active site about 2.5% of the time.

The cGMP sites of both CNBs are accessible to solvent in the auto-inhibited state (***Figure supplement 10***), so cGMP could bind to either of these when its intracellular concentration rises; binding to the high affinity A site may be more likely (***Figure 8***, middle left). Binding of cGMP to CNB-A causes a conformational change that reorients PBC-A, forces CNB-A away from the C-domain, and enhances the tendency of the AI to leave the active site, consistent with the partial activation associated with cGMP binding to the high affinity site binding (Corbin and Doskeland, 1983).

Binding of a second cGMP to the lower affinity, highly selective B site of a kinase monomer (***Figure 8***, middle right) accompanies reorganization of the helical subdomain as seen in the isolated CNB-B domain bound to cGMP (***Figure 5***) (Huang et al., 2014b), weakens R:C interactions at Sites 3 and 4, and releases the CNB-B αA and αB helices from the bottom of the C-domain. With neither CNB bound to the C-domain, the domains diffuse relatively independently and the interdomain linker can sample non-helical conformations, especially around G220, consistent with small angle scattering data for monomeric PKG Iβ showing that cGMP binding to both sites is required for R:C dissociation, elongation, and full activation (Wall et al., 2003). The auto-inhibited state fixes the relative positions of domains within a monomer (***Figure 8***, bottom left), but in activated states these domains may behave in solution like beads on a string.

If each monomer of a leucine zipper-mediated dimer has two bound cGMP, the PBC-A and bent interdomain helices of one monomer can associate with the same regions of the other monomer and assemble into an antiparallel R:R interface (***Figure 4C***) (Kim et al., 2016) that cooperatively stabilizes the activated state (***Figure 8***, bottom right). The R:R interaction explains why the full-length enzyme shows cooperativity but leucine zipper deletion mutants do not (Doskeland et al., 1987; Wall et al., 2003). This assembly helps sustain kinase activation since it occupies much of the R-domain surface that would otherwise be free to associate with the C-domain; it also shields the bound cGMP from solvent, which reduces the rate at which it might be released and degraded by phosphodiesterases. Fixing the CNB domain orientations in this complex may also influence access of the AI sequence to the kinase active site. Dynamic assembly and disassembly of the R:R interface and release and re-binding of cGMP would allow the occupancy of the R:R state to be coupled to the instantaneous cGMP concentration; when cGMP levels fall, the R:R state will be de-populated, and the R-domain will be available to interact with and inhibit the C-domain.

In this model of cGMP-mediated PKG activation (***Figure 8***), the rapid, reversible sampling of states is biased away from the auto-inhibited enzyme default by PBC conformational changes from ‘open’ to ‘closed’ that are coupled to binding of cGMP. The model can be expanded by adding states of the holoenzyme. *R_P_*-cGMPS, which can bind to CNB domains without inducing the ‘closed’ state (Campbell et al., 2017), favors auto-inhibition by stabilizing enzyme states with ‘open’ PBC loops and by competing with cyclic nucleotides that would otherwise bind and induce ‘closed’ PBC loops, explaining why it blocks activation but not basal activity. Because cAMP can bind to the CNBs and induce the closed state (VanSchouwen et al., 2015b), the model predicts that cAMP will act as a partial agonist for the holoenzyme, especially through binding to CNB-A, which discriminates poorly between cGMP and cAMP (Kim et al., 2011). Mutations that favor the PBC ‘closed’ state, or that favor the conformational changes that occur in response to the ‘open’ to ‘closed’ transition, will also favor activation. The activating effect of the CNB-A RQ mutation (Chan et al., 2020; Guo et al., 2013) reflects destabilization of the auto-inhibited state we describe here.

The activation model (***Figure 8***) can explain the substrate-dependence of the PKG Iβ cGMP activation constant (***Figures 1D and Figure supplement 3***) when its states are expanded to include competition between inter- and intra-molecular binding at the active site. Substrate binding pulls the coupled equilibria of ***Figure 8*** to the right. The degree to which the activation constant shifts should depend on the affinity and concentration of the peptide, which determine the bound fraction in competition with binding by the AI sequence (the PKG Iβ sequence makes more favorable contacts than would be expected for the PKG Iα sequence). Substrate-induced shifting of the activation curve rules out models in which substrate binding occurs only after activation; the coupled equilibria imply that every species in ***Figure 8*** with a free active site cleft can reversibly adopt a corresponding substrate-bound state. Co-localization of PKG Iβ with targets through its isotype-specific N-terminal domain (Casteel et al., 2005) may therefore not only influence the choice of substrate but also enhance the degree to which the enzyme is activated.

Although the auto-inhibited state described here (***Figure 8***, bottom left) fixes the relative positions of the CNBs and C-domain, in other states the interdomain linkers may allow a monomer to behave more dynamically, like domain beads on a string. The shorter linker and the distinct leucine zipper and hinge regions of PKG Iα may decrease the opportunities for random diffusion to juxtapose the AI with the active site cleft, resulting in a lower AI effective local concentration at the active cleft for PKG Iα compared to Iβ. This physical rationale would explain why PKG Iα shows minimal substrate-dependence in its cGMP activation constant: substrates will compete better with an AI that more rarely accesses the active site cleft. This rationale may also explain how changes to Iα-specific residues outside the AI impact activation (Ruth et al., 1997). Mutation C43S substantially increases the cGMP activation constant for PKG Iα (Kalyanaraman et al., 2017), which could be explained by the loss of a C43-mediated disulfide in the leucine zipper (Qin et al., 2015) that increases dynamic motions and facilitates AI access to the kinase active site cleft.

This model and our structure provide rationales for how changes in the AI affect Iα and Iβ isoform cGMP activation constants and fractional basal activities by affecting AI contacts with the active site, but they do not provide detailed insights as to how the sequences and structures of the isoform-specific leucine zippers and other elements in the N-terminal domain might affect access to any of the states in the model. The only explicit role for the leucine zippers in the model of ***Figure 8*** is to promote sampling of the R:R state, whose interacting residues are present in both isoforms; how isoform differences might structurally influence access to this state is not known since the leucine zippers were truncated in the construct used for crystallization. Explanations for some of the observed functional differences between the Iα and Iβ isoforms must await more detailed models, and perhaps structures of full-length enzyme isoforms in different states of activation or inhibition.

## Materials and Methods

### Construct design, protein expression and purification of PKG Iβ (71-686)

The sequence encoding human PKG Iβ (residues 71-686) was cloned into pBlueBacHis2A vector with tobacco etch virus (TEV) protease site before coding sequence. The protein was expressed in SF9 cells with multiplicity of infection of 2 for 48 h. Cells were suspended in 200 mM potassium phosphate (pH 7.5), 500 mM NaCl (sodium chloride), 1 mM β-mercaptoethanol (Buffer A) and lysed with a cell disruptor (Constant Systems, Daventry Northants, UK). The supernatant was loaded onto HisTrap^TM^ HP 5 ml column (GE healthcare) and eluted with buffer A containing 300 mM imidazole on an ÄKTA purifier system (GE Healthcare). The his-tag was removed by incubating the eluted protein with TEV protease overnight at 4 °C. A second nickel affinity chromatography step was performed to remove TEV protease and flow-through fractions were collected. The tagless protein was further purified by anion exchange chromatography (Mono Q 10/100 GL, GE Healthcare) in 25 mM potassium phosphate (pH 7.5), 1 mM β-mercaptoethanol) with and without 1 M NaCl. The Mono Q peak fractions were pooled, concentrated and passed through a Hiload 10/300 Superdex 200 column (GE Healthcare) equilibrated with 25 mM MES, 150 mM NaCl, 1 mM tris(2-carboxyethyl)phosphine (TCEP), 2 mM MnCl_2_, and 0.1 mM AMP-PNP.

### Expression and Purification of PKG Iα (79–212)

A DNA construct containing His-tagged human PKG Iα (79–212) R177Q was cloned into pQTEV and transformed into TP2000 *E. coli* cells (Kim et al., 2015b; Roy and Danchin, 1981). Cells were grown at 37°C until OD_600_ of 0.6 and induced with 0.5 mM IPTG. Cells were then grown for an additional 10 hours at 25°C, harvested by centrifugation. Cells were suspended in Buffer A (50 mM potassium phosphate, 500 mM NaCl, 1 mM β-mercaptoethanol (pH 7.5)) and lysed with a cell disruptor (Constant Systems, Daventry Northants, UK)). The protein was purified with BioRad IMAC resin on a BioRad Profinia purification system and eluted with Buffer A containing 300 mM imidazole. The elution fraction protein sample was incubated with TEV protease at 4°C overnight for His-tag removal. A second nickel affinity chromatography step was performed to remove TEV protease and flow-through fractions were collected. Protein was further purified with gel filtration on a Hi-load 16/60 Superdex-75 column (GE Healthcare) in 25 mM Trizma (pH 7.5), 150 mM NaCl, and 1 mM TCEP-HCl.

### Crystallization, Data Collection, Phasing, Model Building, and Refinement

To obtain crystals, PKG Iβ (71-686) protein was concentrated to 20 mg ml^-1^ by using 30 kDa cutoff Amicon Ultra (Millipore) and initial crystal screening was performed by Mosquito Crystal robot (TTP Labtech) with 100 nL protein drop and 100 nL crystallization solution over 70 µL well. Further optimization was performed with 250 nL protein and 250 nL crystallization solution. Crystals were obtained by vapor diffusion method in 120 mM ethylene glycol, 100 mM Bicine (pH 8.5), 20 % w/v PEG 8,000 at room temperature. The crystals were flash cooled in Berkeley, CA, USA). To obtain crystals of PKG Iα (79–212) R177Q, 10 mM cGMP (Aral Biosynthetics) was added to the purified protein, and concentrated to 20 mg ml^-1^ with a 10 kDa cutoff Amicon Ultra (Millipore). Initial crystal screening was performed by Mosquito Crystal robot (TTP Labtech) with 300 nL protein drop and 300 nL crystallization solution over 70 µL well. Further optimization was performed with 2 µL protein and 2 µL crystallization solution over 500 µL well. Crystals were obtained by vapor diffusion method in 6% Tacsimate (pH 4.5), 18% PEG 3350 at 22°C. The R177Q crystals were dipped in cryoprotectant (Paratone-N) and flash cooled in liquid nitrogen. Diffraction data was collected on Beamline 5.0.1 (Advanced Light Source, Berkeley, CA, USA). Diffraction data for both crystals were processed by using CCP4.iMosflm (Battye et al., 2011) and structure determined by Phaser MR using the crystal structure of PKA holoenzyme (PDB ID: 2QCS) and cGMP bound PKG Iβ CNB-A structure (PDB ID: 4Z07) as molecular replacement probes (McCoy et al., 2007). Coot was used for model building of the structures and Phenix.Refine was used for refinement (Afonine et al., 2012; Emsley and Cowtan, 2004). All the figures were generated using PyMOL (Delano -Scientific).

### Small Angle X-ray Scattering Data Collection and Analysis

PKG Iβ (71-686) protein at 3.5 mg ml^-1^ at 20°C was exposed for 3 s at ALS beamline 12.3.1 and SAXS data were collected over the q range 0.01 to 0.39 Å^-1^. Primary data reduction and processing was performed with the ScÅtter pipeline and ATSAS suite. The fitted values for I(0) and R_g_ from Guinier analysis and P(r) analysis agree closely (see ***Table supplement 3***); I(0) is consistent with a monomer and the experimental R_g_ is consistent with the cis-conformation. Ab initio analysis was performed with DAMMIF/DAMMIN, computation of model intensities was performed with the FoXS webserver, and crystal structures were fit into models using SUPCOMB.

### Cloning, protein expression and purification of full-length PKG Iα and Iβ

FLAG-TwinStrep®-tagged hPKG Iα and Iβ were generated by cloning the respective DNA sequence from pFLAG *h*PKG Iα and pFLAG *h*PKG Iβ (Kalyanaraman et al., 2017) using 5’-AAA GCT AGC AGC GAG CTA GAG GAA GAC TTT GCC-3’ and 5’-AAA CTC GAG TTA TTA GAA GTC TAT ATC CCA TCC TGA-3’ as primers for PKG Iα and 5’- GAA GCT AGC ATC CGG GAT TTA CAG TAC G-3’ and 5’- AAA CTC GAG TTA TTA GAA GTC TAT ATC CCA TCC TGA-3’ for PKG Iβ. Both PCR products were subcloned by *Nhe*I and *Xho*I digestion and subsequent ligation into the *Nhe*I and *Xho*I digested pcDNA 3.0 SF TAP plasmid (Gloeckner et al., 2007). The generated plasmids were analyzed by Sanger sequencing. Both constructs were expressed in HEK293-T cells grown to a confluency of ∼80% and transfected with the respective plasmid using polyethylenimine (PEI). Cells were harvested and freeze thawed for purification. Subsequent cell lysis was performed in lysis buffer containing 50 mM Tris (pH 7.4), 150 mM NaCl, 0.5 mM TCEP, protease inhibitor cocktail (Roche) phosphatase inhibitor (Roche) and 0.4% Tween20. Proteins were purified using Strep-Tactin Superflow resin (IBA Lifesciences) employing four washing steps. First, lysis buffer, followed by a high phosphate buffer (366 mM Na_2_HPO_4_, 134 mM NaH_2_PO_4_, 0.5 mM TCEP) and two times Tris buffer (50 mM Tris, 150 mM NaCl, 0.5 mM TCEP). Elution was performed using a Tris buffer containing 2.5 mM desthiobiotin. Subsequently, a buffer exchange with Tris buffer was performed to store the proteins without the desthiobiotin to avoid possible interference with downstream analyses.

### PKG Iα and Iβ in vitro Kinase Assays

Specific kinase activities, activation constants (*K*_act_) and Michaelis-Menten constants (*K*_M_) were determined using an enzyme-coupled spectrophotometric kinase assay as described by Cook et al. (1982). For specific kinase activity, assay buffer (100 mM MOPS, 10 mM MgCl_2_, 1 mM ATP, 1 mM phosphoenolpyruvate, 15.1 U/ml lactate dehydrogenase, 8.4 U/ml pyruvate kinase, 230 µM NADH, 5 mM β-mercaptoethanol) was mixed with the hPKG samples in a glass cuvette and the reaction was started by adding the respective substrate peptide (VASPtide: measurements were performed without cGMP. Maximal kinase activity was determined in presence of a final cGMP concentration of 200 µM. The absorption at 340 nm was monitored for 30-120 s using a double beam photometer (Specord 205, Analytik Jena) and the slope was determined to calculate the specific kinase activity.

The activation constant (*K*_act_) was determined by mixing assay mix with a kinase mix and a dilution series of cGMP in a 384-well plate (Microplate, 384 well, PS, F-bottom, clear, Greiner Bio-One) in a ratio of 2:1:1 and monitoring the absorption at 340 nm for 200-300 s in a microplate reader (CLARIOstar, BMG Labtech). Protein activity was calculated according to the Lambert-Beer law and converted to the observed catalytic activity at the given substrate concentration of 1 mM (*k*_obs_). Values of *k*_obs_ were plotted against the cGMP concentration on a logarithmic scale and *K*_act_ was determined by fitting the data using a sigmoidal dose-response fit with a variable slope. The activation constant is defined as the cGMP concentration at which half-maximal kinase activity occurs.

A similar procedure was followed for the determination of Michaelis-Menten constants (*K*_M_). Here, assay mix, kinase mix with cGMP (final: 50 µM cGMP) and a dilution of VASPtide or Kemptide, respectively, were mixed in a 384 well microplate in a 2:1:1 ratio and processed as described. Calculated kinase activities were plotted against the substrate concentration and fitted to the Michaelis-Menten equation to obtain *K*_M_ and *v*_max_.

### PKG Iβ heterodimer in vitro Kinase Assays

The PKG Iβ complexes were purified from 293T cells co-transfected with Flag-tagged dead and untagged active PKG Iβ expression plasmids, as described (Chan et al., 2020). Purified proteins were diluted in KPEM Buffer (10 mM potassium phosphate (pH 7.0), 1 mM EDTA, and 25 mM mercaptoethanol) and reactions were initiated by adding to reaction mix with or without 10 μM cGMP. Final reactions contained: 40 mM HEPES (pH 7.0), 8 µg Kemptide, 10 mM MgCl_2_, 60 µM ATP, and 0.6 μCi ^32^P-γ-ATP. Reactions were incubated at 30 °C for 1.5 min. and stopped by spotting on P81 phosphocellulose paper. Unincorporated ^32^P-γ-ATP was removed by washing in 4 times 2 liters of 0.452% o-phosphoric acid, and ^32^PO_4_ incorporation was measured by liquid scintillation counting.

### PKG Iβ CNB-A domain NMR

The PKG Iβ 92-227 construct was expressed with an N-terminal His tag, in BL21(DE3) *E.coli* cells grown at 37°C in ^15^N M9 media. Expression was induced with 0.5 mM IPTG at an OD_600_ of 0.8, and the cells were grown for 16 hours at 18 °C. PKG Iβ 92-227 was purified as reported (VanSchouwen et al., 2015b). NMR samples were prepared in 50 mM Tris, pH 7.0, 100 mM NaCl, 1 mM DTT, 0.2%(w/v) NaN_3_. An apo sample was prepared by concentrating the purified PKG to 100 μM, and adding 5% (v/v) D_2_O, PDE, 1 mM ATP and 10 mM MgCl_2_. A cGMP-bound sample was prepared similarly, with the addition of 1 mM cGMP to the apo protein solution. All HSQC spectra were recorded with 16 scans and 1 s recycle delay. The spectra included 128 (t1) and 2048 (t2) complex points with spectral widths of 38.0 ppm and 16.2 ppm for the ^15^N and ^1^H dimensions, respectively. Carrier frequencies of ^1^H and ^15^N were set at the water and central amide region, respectively. All spectra were acquired at 300 K with a Bruker Avance 700 MHz NMR spectrometer equipped with a 5 mm TCI cryoprobe. The spectra were processed with TOPSPIN and analyzed using Sparky (Lee et al., 2015).

### MD Simulation Protocol

All MD simulations were performed using the NAMD 2.12 software (Phillips et al., 2005) on the Shared Hierarchical Academic Research Computing Network (SHARCNET), following a previously described protocol (VanSchouwen and Melacini, 2018), with solvent box dimensions of 49 Å for the wild-type CNB-A domain structures, 50 Å for the RQ-mutant CNB-A domain structure, or 186 Å for the full-length cGMP-free dimer structure. All simulations were executed using 2.1 GHz 32-core Broadwell compute nodes accelerated with two NVIDIA Pascal GPUs per node. The CNB-A domain simulations were each executed for >500 ns at constant temperature and pressure, while the full-length cGMP-free dimer simulation was executed in triplicate for 200 ns at constant temperature and pressure, saving structures every 100000 timesteps (*i.e.* 100.0 ps) for subsequent analysis. A summary of the MD simulations is given in ***Table supplement 3***.

## Acknowledgments

We thank Kim lab members for critical reading of the manuscript. C.K. was funded by NIH grant R01 GM090161. F.W.H. was supported by a grant of the Deutsche Forschungsgemeinschaft (He1818/10-1), the Kassel Graduate school “clocks”, and Federal Ministry of Education and Research, Germany (TargetRD, FKZ: 16GW0270 to F.W.H.). The Berkeley Center for Structural Biology is supported in part by the NIH, the National Institute of General Medical Sciences, and the Howard Hughes Medical Institute. The Advanced Light Source is supported by the Director, Office of Science, Office of Basic Energy Sciences, of the U.S. Department of Energy under contract no. DE-AC02-05CH11231. The SIBYLS beamline (ALS) is supported in part by US DOE program Integrated Diffraction Analysis Technologies and NIH project ALS-ENABLE (P30 GM124169) and a High-End Instrumentation Grant S10OD018483.

## Author Contributions

R.S., P.H., F.W.H., D.E.C., and C.K. designed the experiments addressing the questions in this study. D.E.C. designed the constructs and R.S., L.Q., J.J.K., and C.K. developed purification protocols for crystallization. R.S. obtained crystals of PKG Iβ 71-686 and J.J.K. obtained crystals of PKG Iα 79-212 R177Q. B.S. collected diffraction images. R.S., J.J.K., and C.K. solved the structures and refined the crystallographic models. R.S., K.R.M., J.J.K., and C.K. analyzed structures. P.H. designed and purified full-length Twin-Strep-tagged PKG Iα and Iβ wild type constructs, measured kinase activation constants, basal activity, and fold activation. D.E.C. designed and purified the full-length PKG Iβ wild type and mutant constructs for in vitro kinase assays to test inter-chain communication. M.A. acquired and processed NMR data and B.B. performed MD simulations. G.M. analyzed NMR data and MD simulations. G.K. analyzed SAXS data. D.E.C., K.R.M., P.H., G.M., F.W.H. and C.K. wrote the manuscript and created the figures. All authors commented on the manuscript.

## Supplemental information for

**Figure supplement 1.**
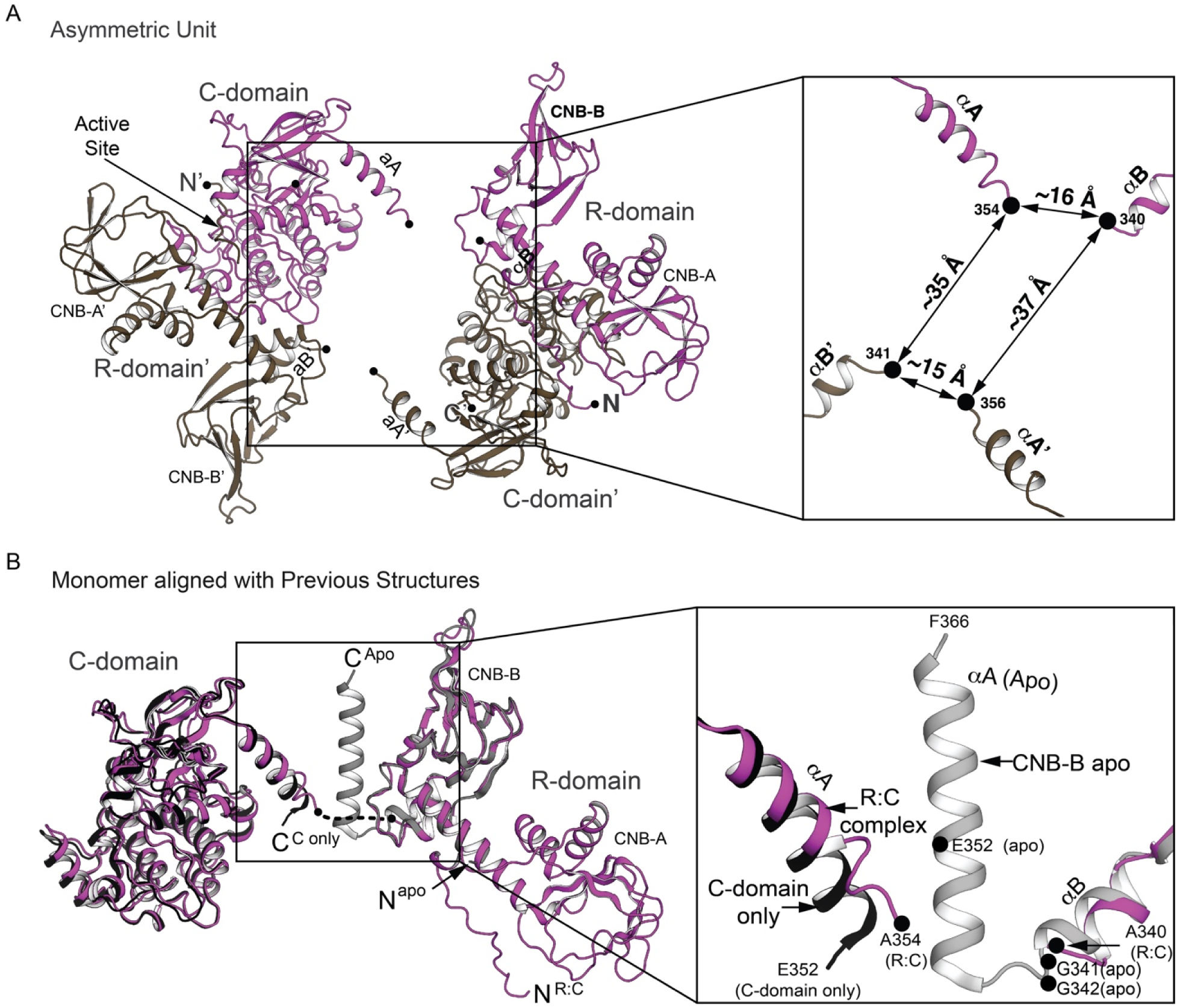
(A) Asymmetric unit. Left: The dimer captured in asymmetric unit with each monomer colored in magenta or dark tan. Right: The zoom-in panel shows the R:C linker at the missing electron density with the distances between the last fitted atoms. (B) The R:C monomer alignment with previous structures. Left: Structural alignment of the R:C monomer with the C-domain bound to N46 (PDB ID: 6C0T) and the CNB-B apo structure (PDB ID: 4KU8). The missing linker region is shown as dotted line. Right: The zoom-in view shows the R:C linker region.

**Figure supplement 2.**
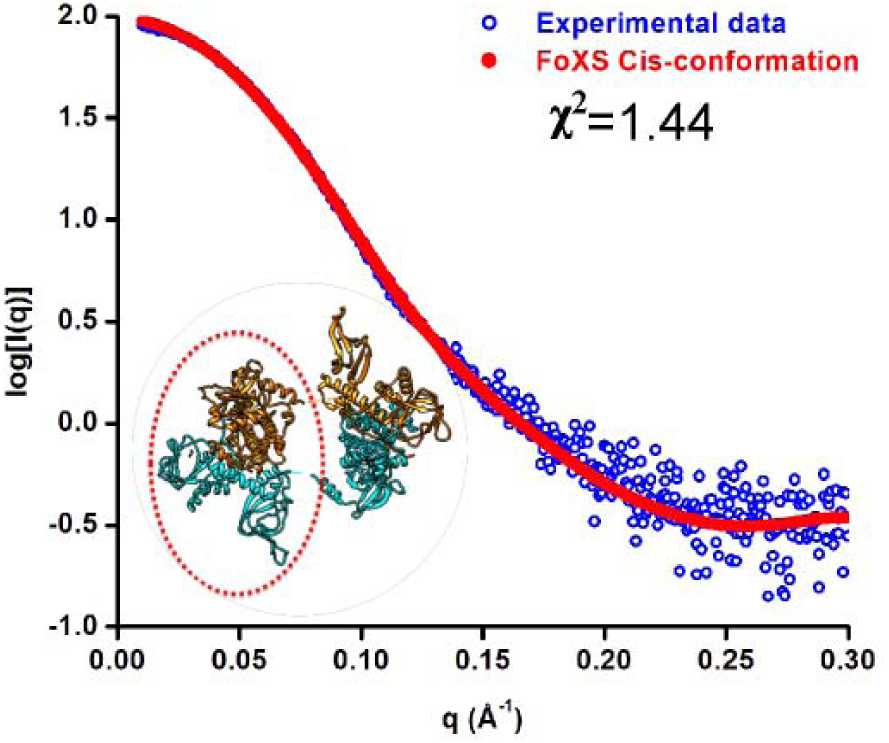
The FoXS webserver analysis shows that the theoretically calculated scattering profile from the cis-conformation (red) matches well with the experimentally observed scattering profile (blue) with a **χ^2 =^** 1.44

**Figure supplement 3.**
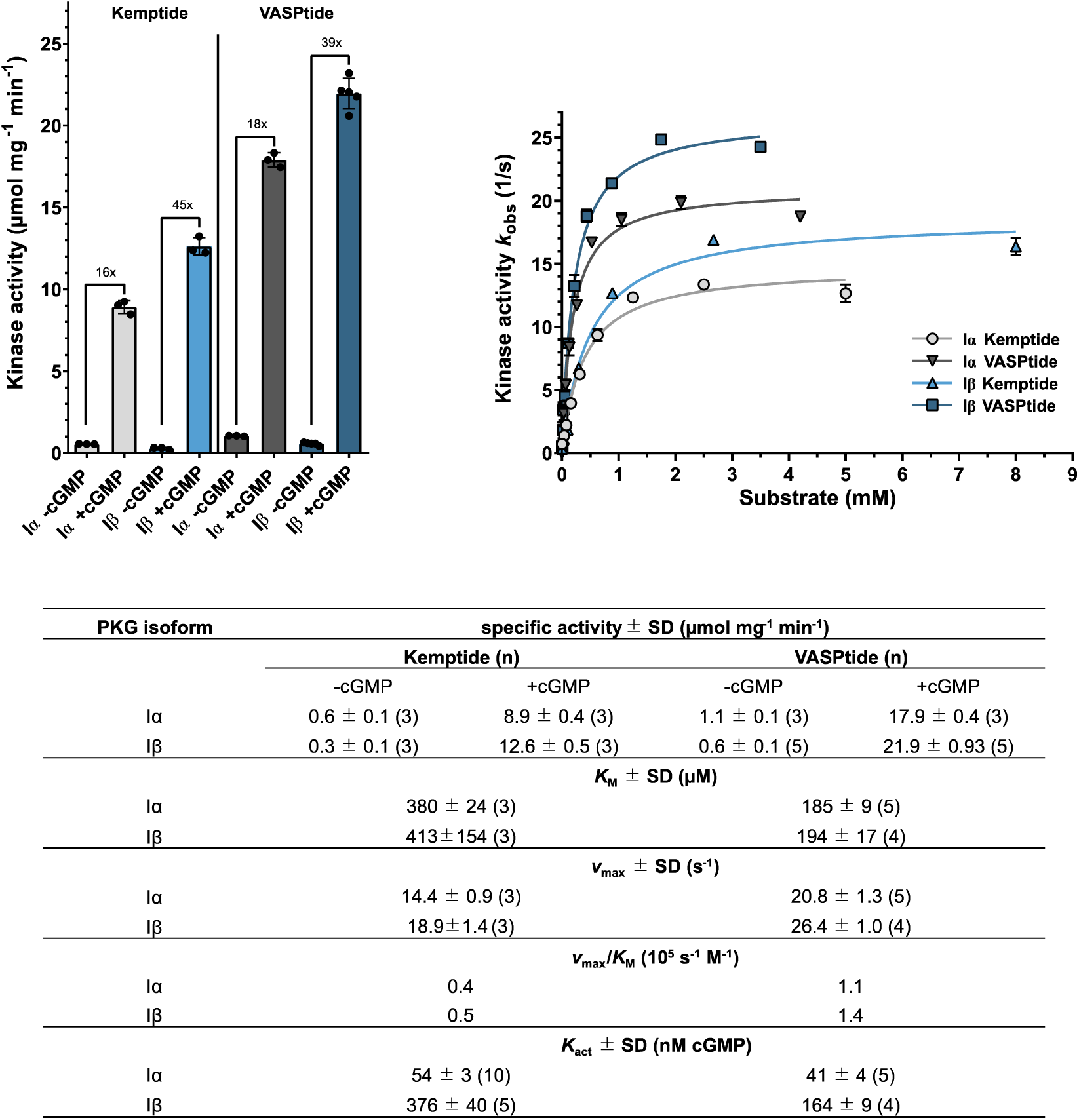
Isoform specific differences between PKG Iα and Iβ. Top left: Specific kinase activities of PKG Iα (grey) and Iβ (blue) in unstimulated (-cGMP) and stimulated (+cGMP) condition using 1 mM of the peptide substrates VASPtide (RRKVSKQE, dark grey and dark blue) or Kemptide (LRRASLG, light grey and light blue). Comparison of both isozymes shows a higher basal activity for PKG Iα and PKG Iβ being more active when saturated with cGMP. Top right: Determination of the Michaelis-Menten constant *K*_M_ and the maximum kinase activity *v*_max_ again show the higher kinase activity of PKG Iβ but most interestingly no difference in the *K*_M_ for the respective substrate is detected. All measurements were performed as spectrophotometric kinase assays according to (Cook et al., 1982). Data points for Michaelis-Menten kinetics are depicted as mean of duplicates with error bars indicating the standard deviation (SD). All values are given as mean of *n* measurements ± SD.

---

**Figure supplement 4.**
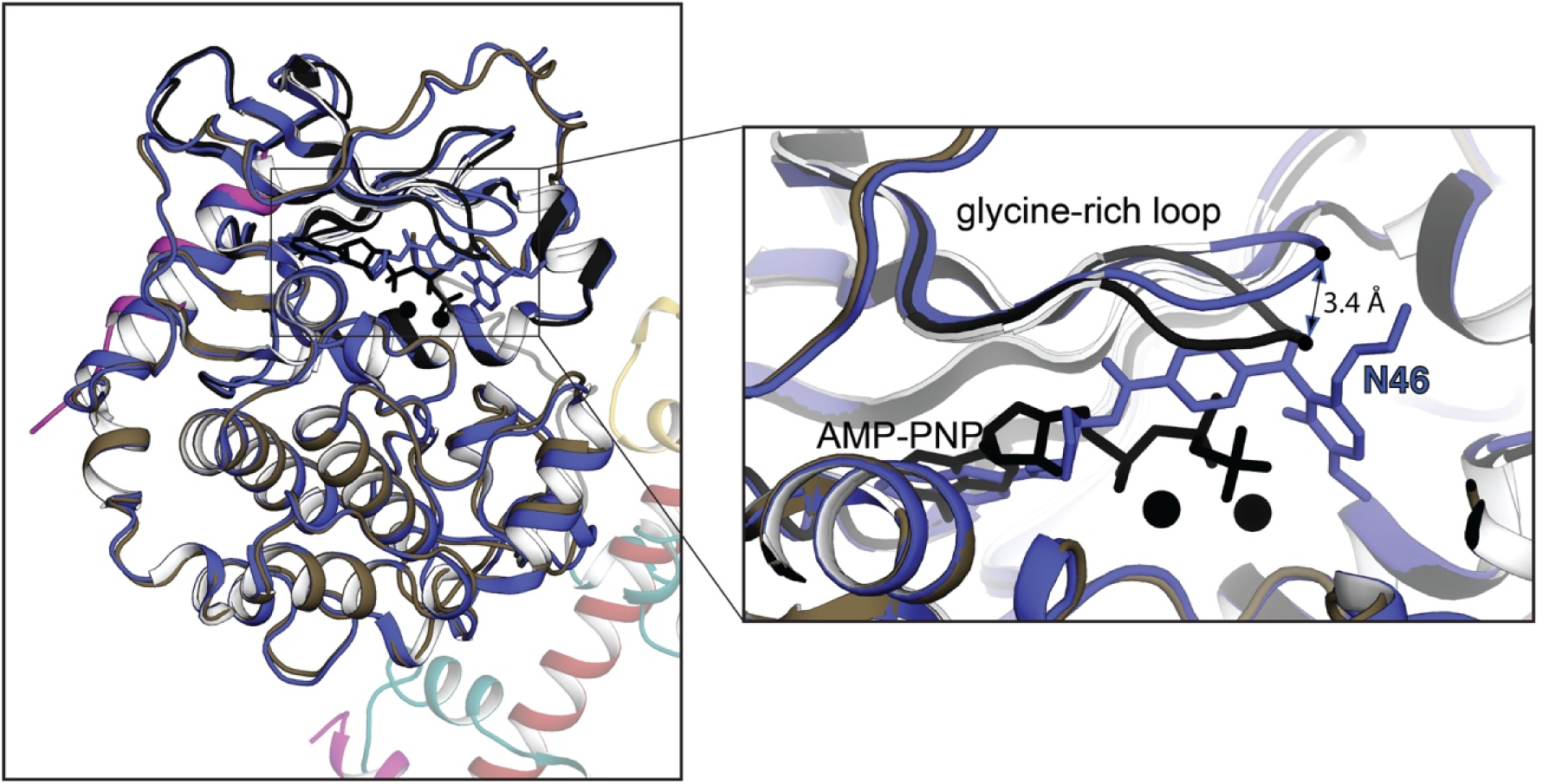
Structural alignment of auto-inhibited PKG Iβ monomer with the isolated PKG I C-domain bound to N46 (PDB ID: 6C0T). The isolated C-domain is in blue and the auto-inhibited R-C complex is in the same color theme as Figure 1B. The zoom-in view on the right highlights the structural differences at the glycine rich loop.

**Figure supplement 5.**
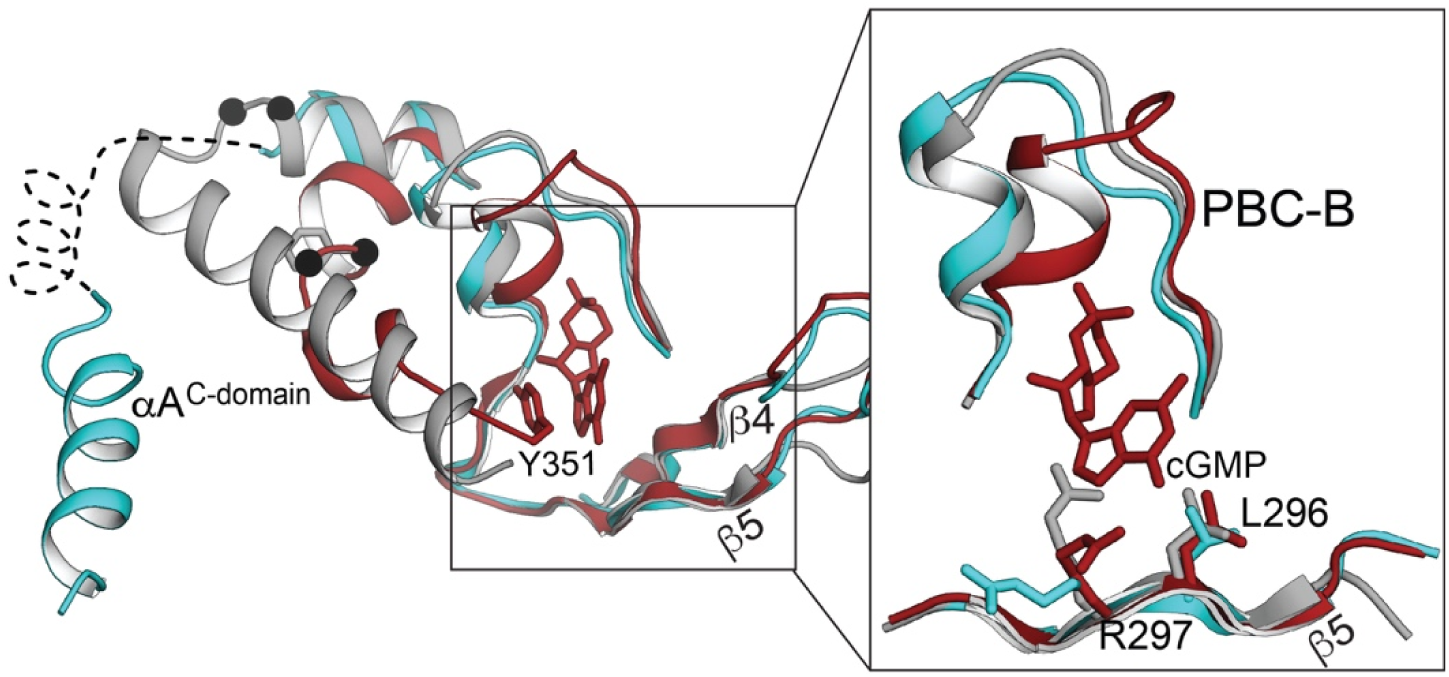
Superposition of the PKG Iβ CNB-B domain in the auto-inhibited state (teal, PDB ID: 7LV3) with the isolated CNB-B domain apo state (red, PDB ID: 4KU8) and cGMP-bound state (gray, PDB ID: 4Z07). Left: Only PBC, β4, β5 and interdomain helices are shown. Right: Zoomed-in view showing PBC and B helix only. Key cGMP-interacting residues of β5 are shown in stick.

**Figure supplement 6.**
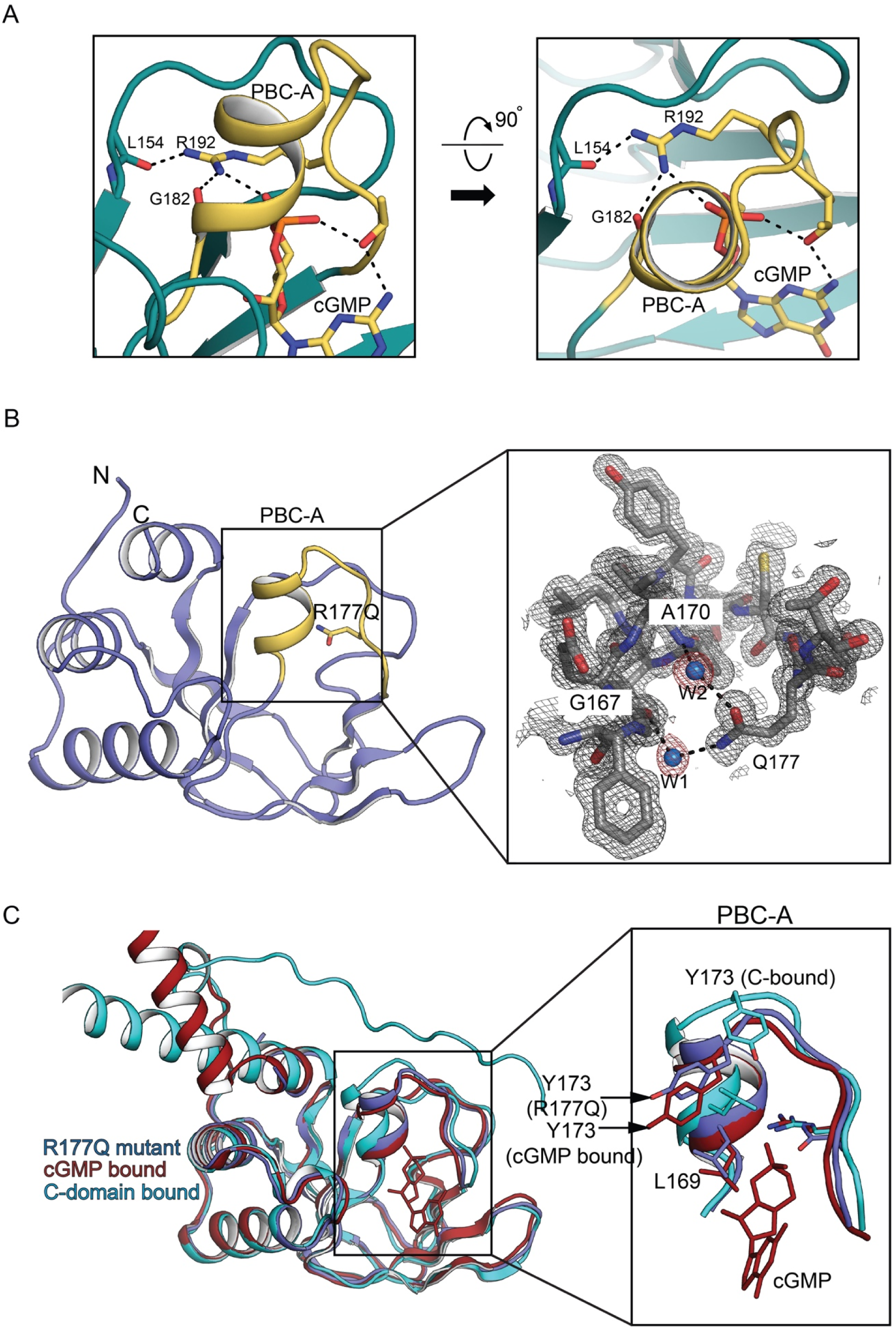
Structural basis of the constitutive activation in TAAD causing mutant. (A) PBC-A R192 interactions in wild type PKG Iβ (PDB ID: 3OD0) (B) Left: Overall structure of PKG Iα CNB-A R177Q (PDB ID: 7MBJ) determined at 1.26 Å. Structure is shown in cartoon with PBC in yellow and R177Q in stick. Right: Zoomed-in view of PBC-A with electron density (2*F_o_-F_c_* at σ =1.0). PBC residues interacting with ordered water molecules and side chain of Q177 are shown in sticks. Ordered water molecules are shown as blue spheres. (C) Structural alignment of CNB-A R177Q with wild type CNB-A bound to cGMP and CNB-A contacting the C-domain (R:C complex). Left: R177Q mutant, cGMP-bound and C-bound CNB-A conformations. The cGMP-bound CNB-A is colored in red, C-domain bound in teal, and R177Q mutant in violet. Right: Zoomed in view only showing PBC. The key R:C interface residues are shown in sticks.

**Figure supplement 7.**
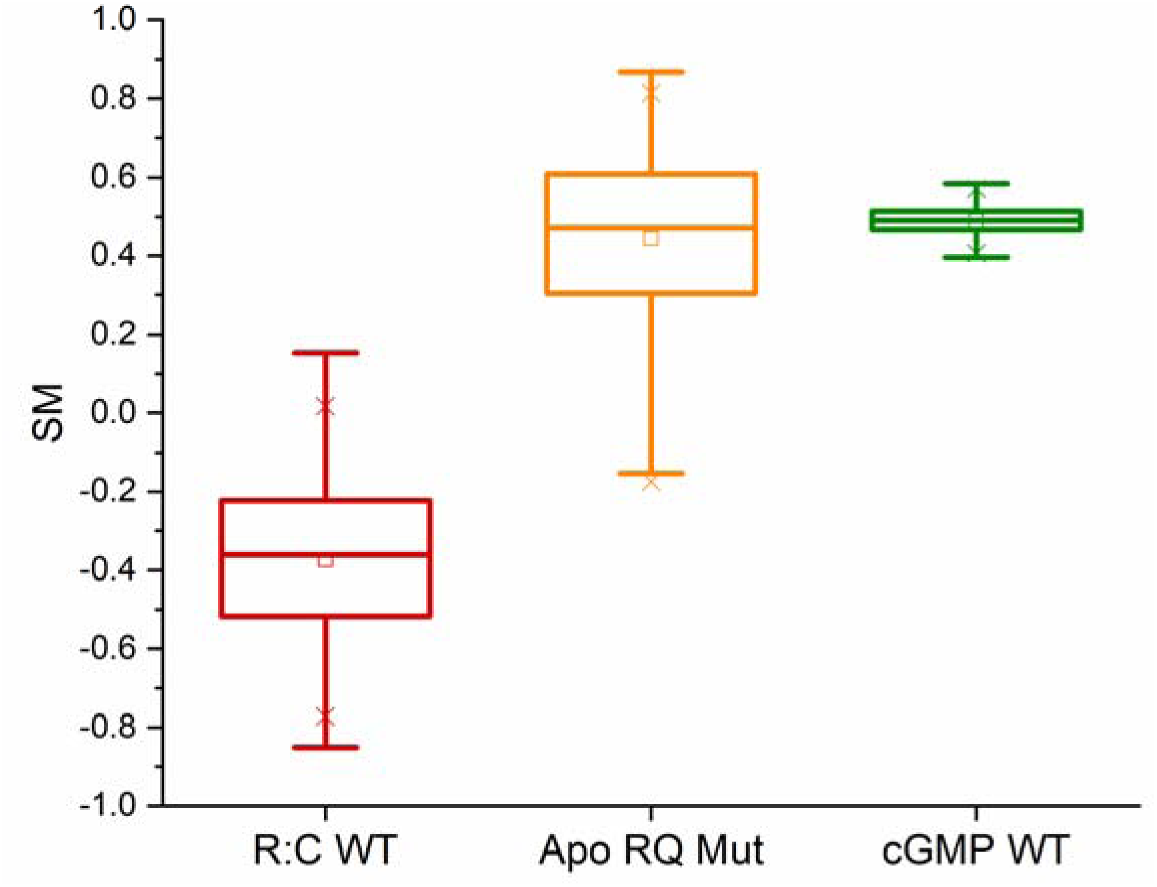
MD simulations. CNB-A domain PBC similarity measures (SM) computed from the three replicate simulations of cGMP-free dimeric PKG Iβ (“R:C WT”), and from the simulations of the isolated CNB-A domain (“Apo RQ Mut” and “cGMP WT”), using the CNB-A domain PBC from the “4Z07” and cGMP-free dimer X-ray structures as active- and inactive-state reference structures, respectively. The statistics reported in each boxplot are as follows: the middle, bottom and top lines of the central box represent the median, 25^th^ percentile and 75^th^ percentile of the data set, respectively; the whiskers represent additional data falling within 1.5*IQR above the 75^th^ percentile or below the 25^th^ percentile (where IQR is the difference between the 75^th^ and 25^th^ percentiles); the “□” symbol represents the mean of the data set; and the two “×” symbols represent the 1^st^ and 99^th^ percentiles of the data set.

**Figure supplement 8.**
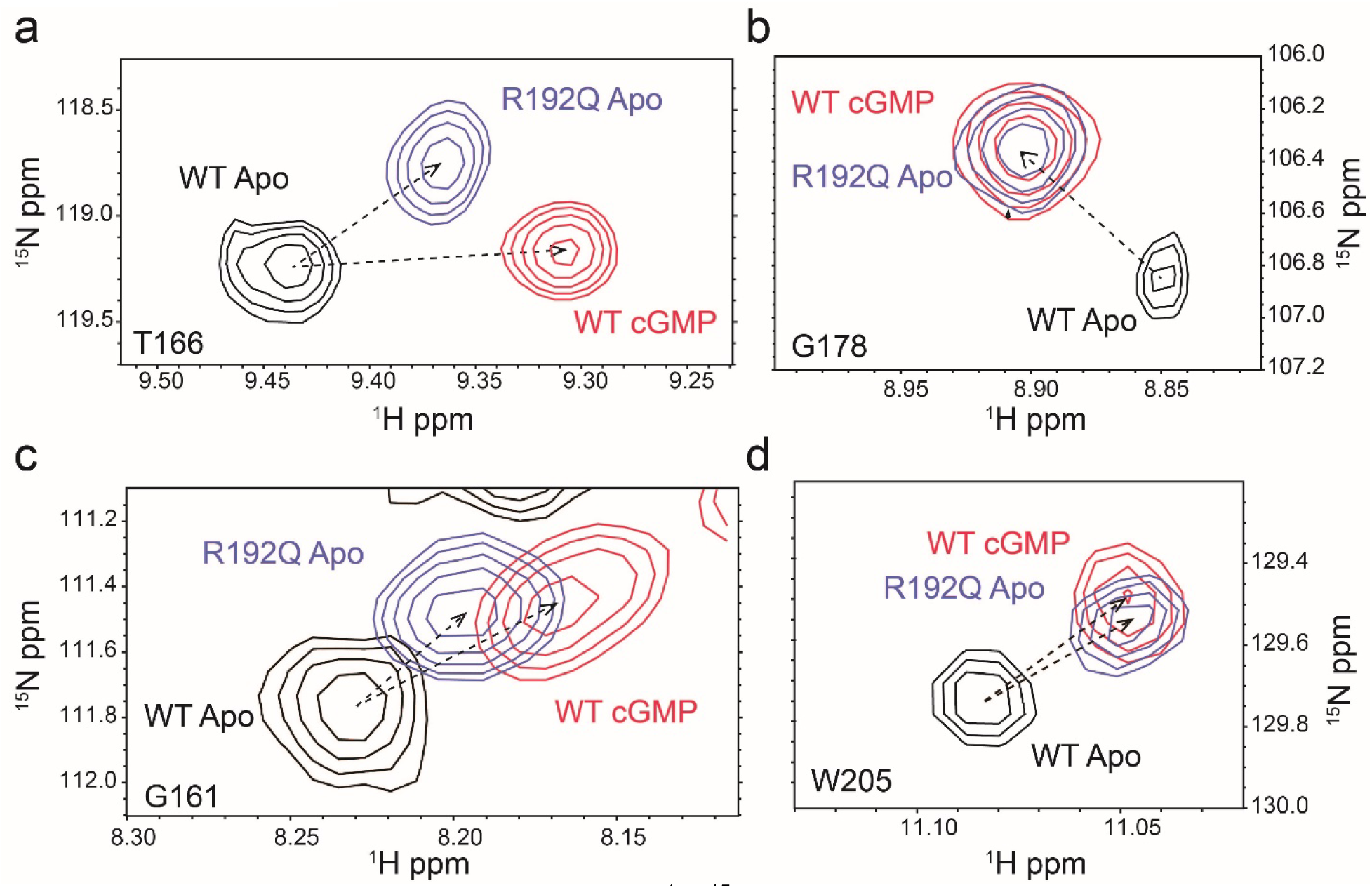
(a - d), representative^1^H,^15^N HSQC cross-peaks of the apo wt (black), cGMP-bound wt (red) and apo R192Q (purple) PKG1b (92-227). For W205 in panel (d), the side chain peak was selected.

**Figure supplement 9.**
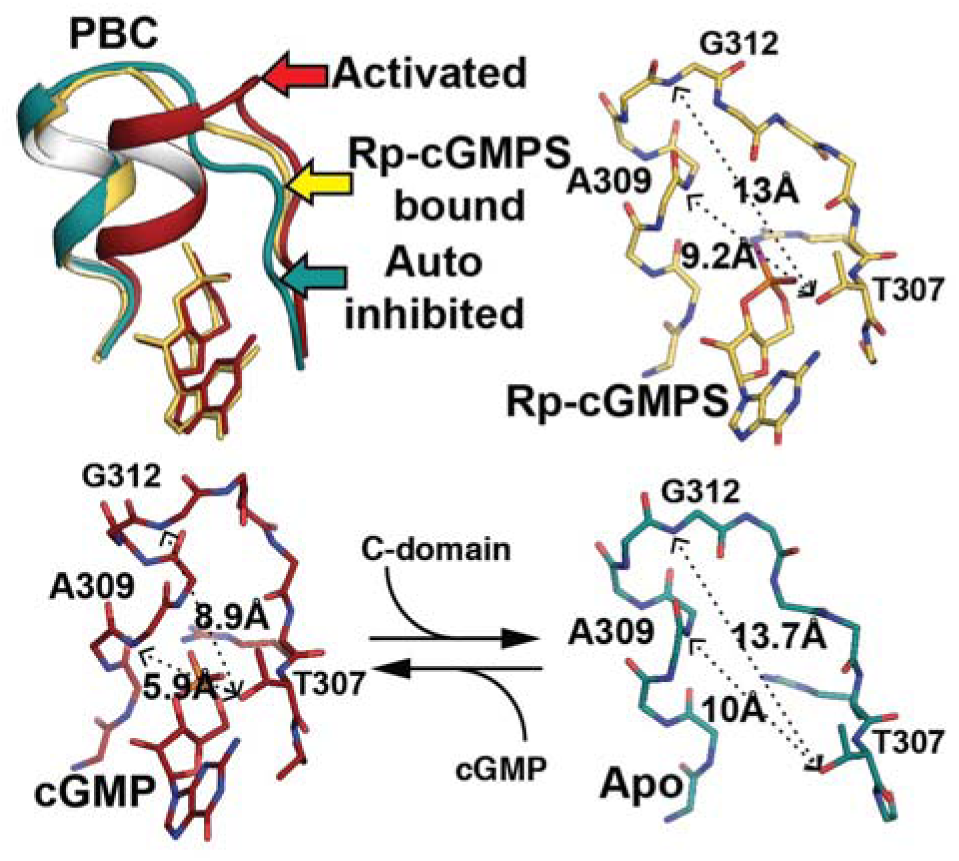
A conformation induced by cGMP is prevented by *R*_P_-cGMPS. Our structure of auto-inhibited PKG I suggests that the first steps in activation are phosphate binding cassette re-ordering and closing of the CNB-B pocket on bound cGMP. Aligning the activated, auto-inhibited, and *R*_P_-cGMPS-bound CNB-B states (top left) shows that the protein backbone closes around cGMP, bringing T307 closer to A309 and G312 (bottom left) than in the auto-inhibited state (bottom right). The *R*_P_-cGMPS-bound state (top right) remains almost as open as the auto-inhibited state.

**Figure supplement 10.**
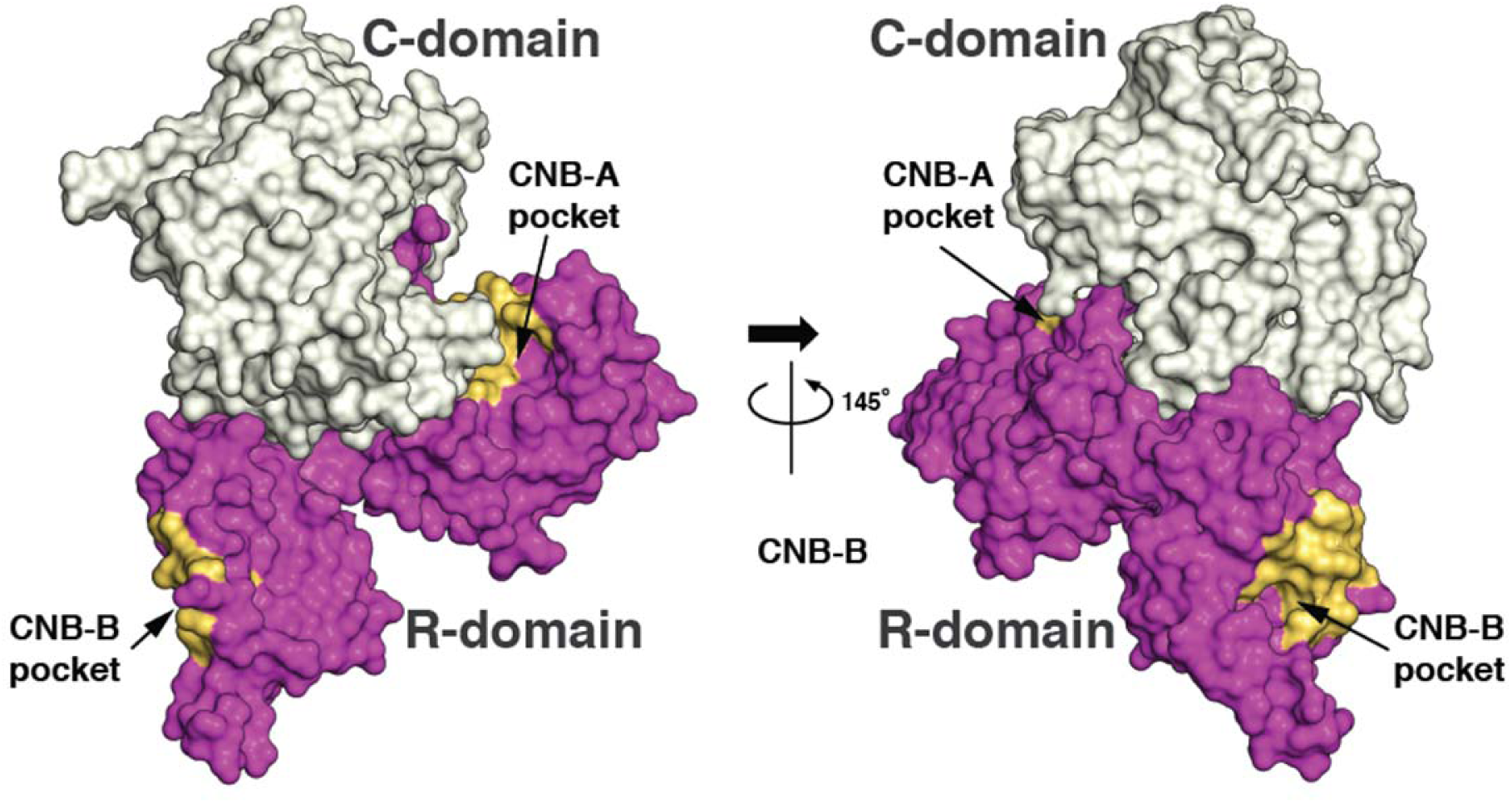
Solvent accessibility of cGMP binding pockets. Surface view of the auto-inhibited monomer showing R-, C-domains and R-subdomain pockets (CNB-A and CNB-B). Both CNB-A and -B pockets are exposed to solvent. The C-domain is shown in grey, R-domain in magenta, CNB-A and CNB-B cGMP pockets in yellow.

**Supplement Table 1.** Data collection and refinement statistics.

| Protein | PKG Iβ 71-686 | PKG Iα 79-212 R177Q | *Information for the highest resolution on shell is shown in parenthesis. |
| --- | --- | --- | --- |
| **Data collection** |  |  |  |
| Wavelength (Å) | 0.97931 | 0.97931 |  |
| Space group | $P2_12_12_1$ | $P1$ | |
| Cell dimensions |  |  |  |
| *a*, *b*, *c* (Å) | 99.8, 112.8, 150.7 | 39.2, 41.9, 48.3 |  |
| $\alpha$, $\beta$, $\gamma$ (°) | 90, 90, 90 | 94.7, 113.7, 114.0 | |
| Resolution (Å) | 47.7 - 2.41 (2.50 - 2.41)* | 36.54 - 1.26 (1.30-1.26)* |  |
| $R_{\text{merge}}$ | 0.1115 (4.83) | 0.06 (0.175) | |
| *I*/σ*I* | 20.92 (0.84) | 25.4 (6.56) | |
| Completeness (%) | 99.95 (99.86) | 93.6 (89.26) |  |
| Redundancy | 13.4 (13.3) | 3.7 (2.8) |  |
| **Refinement** |  |  |  |
| No. reflections | 66392 (6568) | 62350 (3133) |  |
| $R_{\text{work}}/R_{\text{free}}$ (%) | 19.18/23.80 | 17.76/19.69 | |
| No. atoms |  |  |  |
| Proteins | 9208 | 2034 |  |
| Ligand | 110 | 0 |  |
| Water | 87 | 361 |  |
| B-factors |  |  |  |
| Overall | 84.27 | 23.17 |  |
| Protein | 84.53 | 20.91 |  |
| Ligand | 79.41 |  |  |
| Water | 63.0 | 35.90 |  |
| R.m.s. deviations |  |  |  |
| Bond lengths (Å) | 0.009 | 0.009 |  |
| Bond angles (°) | 1.10 | 1.18 |  |
| PDB Code | 7LV3 | 7MBJ |  |

\*intensities were excluded from refinement for cross validation purposes.

**Supplement Table 2.**
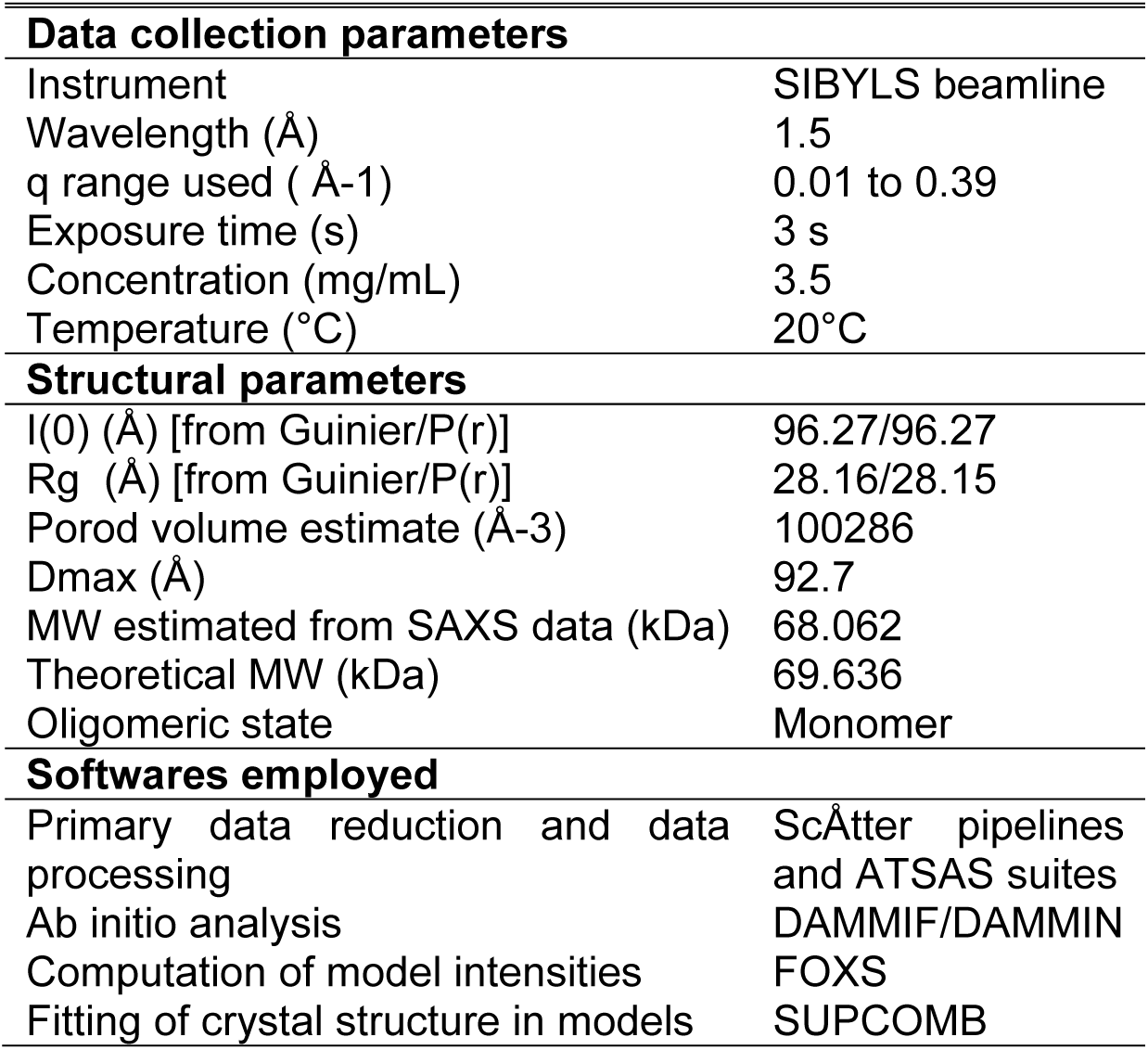
SAXS Data Collection and Scattering-Derived Parameters for PKG Iβ 71-686.

| **Data collection parameters** |  |
| --- | --- |
| Instrument | SIBYLS beamline |
| Wavelength (Å) | 1.5 |
| q range used ( Å<sup>-1</sup>) | 0.01 to 0.39 |
| Exposure time (s) | 3 s |
| Concentration (mg/mL) | 3.5 |
| Temperature (°C) | 20°C |
| **Structural parameters** |  |
| I(0) (Å) [from Guinier/P(r)] | 96.27/96.27 |
| R<sub>g</sub> (Å) [from Guinier/P(r)] | 28.16/28.15 |
| Porod volume estimate (Å<sup>-3</sup>) | 100286 |
| D<sub>max</sub> (Å) | 92.7 |
| MW estimated from SAXS data (kDa) | 68.062 |
| Theoretical MW (kDa) | 69.636 |
| Oligomeric state | Monomer |
| **Softwares employed** |  |
| Primary data reduction and data processing | ScÅtter pipelines and ATSAS suites |
| Ab initio analysis | DAMMIF/DAMMIN |
| Computation of model intensities | FOXS |
| Fitting of crystal structure in models | SUPCOMB |

**Supplement Table 3.** Summary of the MD Simulations Performed for PKG Iβ.

| **Simulated Construct** | **Starting Structure** | **Simulation Length per Replicate (ns)** | **Number of Simulation Replicates Executed** | **Number of CPUs Used** |
| --- | --- | --- | --- | --- |
| Apo Wild-Type CNB-A Domain | Chain A from PDB ID "3OD0" | 579.7 | 1 | 32 |
| cGMP-Bound Wild-Type CNB-A | PDB ID :3OD0 | 591.3 | 1 | 32 |
| Apo RQ-Mutant CNB-A | PKG Iα 79-212 R177Q | 652.3 | 1 | 32 |
| PKG Iβ 71-686 | PKG Iβ 71-686 | 200.0 | 3 | 32 |

## References

Afonine, P.V., Grosse-Kunstleve, R.W., Echols, N., Headd, J.J., Moriarty, N.W., Mustyakimov, M., Terwilliger, T.C., Urzhumtsev, A., Zwart, P., and Adams, P.D. (2012). Towards automated crystallographic structure refinement with phenix.refine. Acta Crystallogr D Biol Crystallogr 68, 352–367.

Akimoto, M., McNicholl, E.T., Ramkissoon, A., Moleschi, K., Taylor, S.S., and Melacini, G. (2015). Mapping the Free Energy Landscape of PKA Inhibition and Activation: A Double-Conformational Selection Model for the Tandem cAMP-Binding Domains of PKA RIalpha. PLoS Biol 13, e1002305.

Alverdi, V., Mazon, H., Versluis, C., Hemrika, W., Esposito, G., van den Heuvel, R., Scholten, A., and Heck, A.J. (2008). cGMP-binding prepares PKG for substrate binding by disclosing the C-terminal domain. J Mol Biol 375, 1380–1393.

Battye, T.G., Kontogiannis, L., Johnson, O., Powell, H.R., and Leslie, A.G. (2011). iMOSFLM: a new graphical interface for diffraction-image processing with MOSFLM. Acta Crystallogr D Biol Crystallogr 67, 271–281.

Berman, H.M., Ten Eyck, L.F., Goodsell, D.S., Haste, N.M., Kornev, A., and Taylor, S.S. (2005). The cAMP binding domain: an ancient signaling module. Proceedings of the National Academy of Sciences of the United States of America 102, 45–50.

Boettcher, A.J., Wu, J., Kim, C., Yang, J., Bruystens, J., Cheung, N., Pennypacker, J.K., Blumenthal, D.A., Kornev, A.P., and Taylor, S.S. (2011). Realizing the allosteric potential of the tetrameric protein kinase A RIalpha holoenzyme. Structure 19, 265–276.

Browning, D.D., Kwon, I.K., and Wang, R. (2010). cGMP-dependent protein kinases as potential targets for colon cancer prevention and treatment. Future Med Chem 2, 65–80.

Busch, J.L., Bessay, E.P., Francis, S.H., and Corbin, J.D. (2002). A conserved serine juxtaposed to the pseudosubstrate site of type I cGMP-dependent protein kinase contributes strongly to autoinhibition and lower cGMP affinity. J Biol Chem 277, 34048–34054.

Byun, J.A., Akimoto, M., VanSchouwen, B., Lazarou, T.S., Taylor, S.S., and Melacini, G. (2020). Allosteric pluripotency as revealed by protein kinase A. Sci Adv 6, eabb1250.

Campbell, J.C., VanSchouwen, B., Lorenz, R., Sankaran, B., Herberg, F.W., Melacini, G., and Kim, C. (2017). Crystal structure of cGMP-dependent protein kinase Ibeta cyclic nucleotide-binding-B domain : Rp-cGMPS complex reveals an apo-like, inactive conformation. FEBS Lett 591, 221–230.

Casteel, D.E., Boss, G.R., and Pilz, R.B. (2005). Identification of the interface between cGMP-dependent protein kinase Ibeta and its interaction partners TFII-I and IRAG reveals a common interaction motif. J Biol Chem 280, 38211–38218.

Casteel, D.E., Smith-Nguyen, E.V., Sankaran, B., Roh, S.H., Pilz, R.B., and Kim, C. (2010). A crystal structure of the cyclic GMP-dependent protein kinase I{beta} dimerization/docking domain reveals molecular details of isoform-specific anchoring. J Biol Chem 285, 32684–32688.

Chan, M.H., Aminzai, S., Hu, T., Taran, A., Li, S., Kim, C., Pilz, R.B., and Casteel, D.E. (2020). A substitution in cGMP-dependent protein kinase 1 associated with aortic disease induces an active conformation in the absence of cGMP. J Biol Chem 295, 10394–10405.

Corbin, J.D., and Doskeland, S.O. (1983). Studies of two different intrachain cGMP-binding sites of cGMP-dependent protein kinase. J Biol Chem 258, 11391–11397.

Das, R., Chowdhury, S., Mazhab-Jafari, M.T., Sildas, S., Selvaratnam, R., and Melacini, G. (2009). Dynamically driven ligand selectivity in cyclic nucleotide binding domains. J Biol Chem 284, 23682–23696.

Doskeland, S.O., Vintermyr, O.K., Corbin, J.D., and Ogreid, D. (1987). Studies on the interactions between the cyclic nucleotide-binding sites of cGMP-dependent protein kinase. J Biol Chem 262, 3534–3540.

Emsley, P., and Cowtan, K. (2004). Coot: model-building tools for molecular graphics. Acta Crystallogr D Biol Crystallogr 60, 2126–2132.

Feil, R., Kellermann, J., and Hofmann, F. (1995). Functional cGMP-dependent protein kinase is phosphorylated in its catalytic domain at threonine-516. Biochemistry 34, 13152–13158.

Feil, R., Lohmann, S.M., de Jonge, H., Walter, U., and Hofmann, F. (2003). Cyclic GMP-dependent protein kinases and the cardiovascular system: insights from genetically modified mice. Circ Res 93, 907–916.

Francis, S.H., Busch, J.L., Corbin, J.D., and Sibley, D. (2010). cGMP-dependent protein kinases and cGMP phosphodiesterases in nitric oxide and cGMP action. Pharmacol Rev 62, 525–563.

Francis, S.H., and Corbin, J.D. (1994). Structure and Function of Cyclic Nucleotide-Dependent Protein Kinases. Annual Review of Physiology 56, 237–272.

Francis, S.H., Smith, J.A., Colbran, J.L., Grimes, K., Walsh, K.A., Kumar, S., and Corbin, J.D. (1996). Arginine 75 in the pseudosubstrate sequence of type Ibeta cGMP-dependent protein kinase is critical for autoinhibition, although autophosphorylated serine 63 is outside this sequence. J Biol Chem 271, 20748–20755.

Gronenborn, A.M. (2009). Protein acrobatics in pairs--dimerization via domain swapping. Curr Opin Struct Biol 19, 39–49.

Guo, D.C., Regalado, E., Casteel, D.E., Santos-Cortez, R.L., Gong, L., Kim, J.J., Dyack, S., Horne, S.G., Chang, G., Jondeau, G., et al. (2013). Recurrent gain-of-function mutation in PRKG1 causes thoracic aortic aneurysms and acute aortic dissections. Am J Hum Genet 93, 398–404.

Hammond, J., and Balligand, J.L. (2012). Nitric oxide synthase and cyclic GMP signaling in cardiac myocytes: from contractility to remodeling. J Mol Cell Cardiol 52, 330–340.

Hofmann, F., Bernhard, D., Lukowski, R., and Weinmeister, P. (2009). cGMP regulated protein kinases (cGK). Handbook of experimental pharmacology, 137–162.

Huang, G.Y., Gerlits, O.O., Blakeley, M.P., Sankaran, B., Kovalevsky, A.Y., and Kim, C. (2014a). Neutron diffraction reveals hydrogen bonds critical for cGMP-selective activation: insights for cGMP-dependent protein kinase agonist design. Biochemistry 53, 6725–6727.

Huang, G.Y., Kim, J.J., Reger, A.S., Lorenz, R., Moon, E.W., Zhao, C., Casteel, D.E., Bertinetti, D., Vanschouwen, B., Selvaratnam, R., et al. (2014b). Structural basis for cyclic-nucleotide selectivity and cGMP-selective activation of PKG I. Structure 22, 116–124.

Johnson, D.A., Akamine, P., Radzio-Andzelm, E., Madhusudan, M., and Taylor, S.S. (2001). Dynamics of cAMP-dependent protein kinase. Chem Rev 101, 2243–2270.

Kalyanaraman, H., Schall, N., and Pilz, R.B. (2018). Nitric oxide and cyclic GMP functions in bone. Nitric Oxide 76, 62–70.

Kalyanaraman, H., Zhuang, S., Pilz, R.B., and Casteel, D.E. (2017). The activity of cGMP-dependent protein kinase Ialpha is not directly regulated by oxidation-induced disulfide formation at cysteine 43. J Biol Chem 292, 8262–8268.

Kannan, N., Haste, N., Taylor, S.S., and Neuwald, A.F. (2007). The hallmark of AGC kinase functional divergence is its C-terminal tail, a cis-acting regulatory module. Proceedings of the National Academy of Sciences of the United States of America 104, 1272–1277.

Kim, C., Cheng, C.Y., Saldanha, S.A., and Taylor, S.S. (2007). PKA-I holoenzyme structure reveals a mechanism for cAMP-dependent activation. Cell 130, 1032–1043.

Kim, C., and Sharma, R. (2021). Cyclic nucleotide selectivity of protein kinase G isozymes. Protein Sci 30, 316–327.

Kim, C., Xuong, N.H., and Taylor, S.S. (2005). Crystal structure of a complex between the catalytic and regulatory (RIalpha) subunits of PKA. Science 307, 690–696.

Kim, J.J., Casteel, D.E., Huang, G., Kwon, T.H., Ren, R.K., Zwart, P., Headd, J.J., Brown, N.G., Chow, D.C., Palzkill, T., et al. (2011). Co-crystal structures of PKG Ibeta (92-227) with cGMP and cAMP reveal the molecular details of cyclic-nucleotide binding. PLoS One 6, e18413.

Kim, J.J., Lorenz, R., Arold, S.T., Reger, A.S., Sankaran, B., Casteel, D.E., Herberg, F.W., and Kim, C. (2016). Crystal Structure of PKG I:cGMP Complex Reveals a cGMP-Mediated Dimeric Interface that Facilitates cGMP-Induced Activation. Structure 24, 710–720.

Klinger, J.R., and Kadowitz, P.J. (2017). The Nitric Oxide Pathway in Pulmonary Vascular Disease. Am J Cardiol 120, S71–S79.

Kornev, A.P., Taylor, S.S., and Ten Eyck, L.F. (2008). A generalized allosteric mechanism for cis-regulated cyclic nucleotide binding domains. PLoS Comput Biol 4, e1000056.

Krissinel, E., and Henrick, K. (2007). Inference of macromolecular assemblies from crystalline state. J Mol Biol 372, 774–797.

Lee, W., Tonelli, M., and Markley, J.L. (2015). NMRFAM-SPARKY: enhanced software for biomolecular NMR spectroscopy. Bioinformatics 31, 1325–1327.

Lu, T.W., Wu, J., Aoto, P.C., Weng, J.H., Ahuja, L.G., Sun, N., Cheng, C.Y., Zhang, P., and Taylor, S.S. (2019). Two PKA RIalpha holoenzyme states define ATP as an isoform-specific orthosteric inhibitor that competes with the allosteric activator, cAMP. Proceedings of the National Academy of Sciences of the United States of America 116, 16347–16356.

Luo, C., Kuner, T., and Kuner, R. (2014). Synaptic plasticity in pathological pain. Trends Neurosci 37, 343–355.

Moleschi, K.J., Akimoto, M., and Melacini, G. (2015). Measurement of State-Specific Association Constants in Allosteric Sensors through Molecular Stapling and NMR. J Am Chem Soc 137, 10777–10785.

Osborne, B.W., Wu, J., McFarland, C.J., Nickl, C.K., Sankaran, B., Casteel, D.E., Woods, V.L., Jr., Kornev, A.P., Taylor, S.S., and Dostmann, W.R. (2011). Crystal structure of cGMP-dependent protein kinase reveals novel site of interchain communication. Structure 19, 1317–1327.

Pfeifer, A., Aszodi, A., Seidler, U., Ruth, P., Hofmann, F., and Fassler, R. (1996). Intestinal secretory defects and dwarfism in mice lacking cGMP-dependent protein kinase II. Science 274, 2082–2086.

Phillips, J.C., Braun, R., Wang, W., Gumbart, J., Tajkhorshid, E., Villa, E., Chipot, C., Skeel, R.D., Kale, L., and Schulten, K. (2005). Scalable molecular dynamics with NAMD. J Comput Chem 26, 1781–1802.

Qin, L., Reger, A.S., Guo, E., Yang, M.P., Zwart, P., Casteel, D.E., and Kim, C. (2015). Structures of cGMP-Dependent Protein Kinase (PKG) Ialpha Leucine Zippers Reveal an Interchain Disulfide Bond Important for Dimer Stability. Biochemistry 54, 4419–4422.

Qin, L., Sankaran, B., Aminzai, S., Casteel, D.E., and Kim, C. (2018). Structural basis for selective inhibition of human PKG Ialpha by the balanol-like compound N46. J Biol Chem 293, 10985–10992.

Rangaswami, H., Marathe, N., Zhuang, S., Chen, Y., Yeh, J.-C., Frangos, J.A., Boss, G.R., and Pilz, R.B. (2009). Type II cGMP-dependent Protein Kinase Mediates Osteoblast Mechanotransduction. The Journal of Biological Chemistry 284, 14796–14808.

Reger, A.S., Yang, M.P., Koide-Yoshida, S., Guo, E., Mehta, S., Yuasa, K., Liu, A., Casteel, D.E., and Kim, C. (2014). Crystal structure of the cGMP-dependent protein kinase II leucine zipper and Rab11b protein complex reveals molecular details of G-kinase-specific interactions. J Biol Chem 289, 25393–25403.

Rehmann, H., Wittinghofer, A., and Bos, J.L. (2007). Capturing cyclic nucleotides in action: snapshots from crystallographic studies. Nat Rev Mol Cell Biol 8, 63–73.

Richie-Jannetta, R., Francis, S.H., and Corbin, J.D. (2003). Dimerization of cGMP-dependent protein kinase Ibeta is mediated by an extensive amino-terminal leucine zipper motif, and dimerization modulates enzyme function. J Biol Chem 278, 50070–50079.

Ruth, P., Pfeifer, A., Kamm, S., Klatt, P., Dostmann, W.R., and Hofmann, F. (1997). Identification of the amino acid sequences responsible for high affinity activation of cGMP kinase Ialpha. J Biol Chem 272, 10522–10528.

Schlossmann, J., Ammendola, A., Ashman, K., Zong, X., Huber, A., Neubauer, G., Wang, G.X., Allescher, H.D., Korth, M., Wilm, M., et al. (2000). Regulation of intracellular calcium by a signalling complex of IRAG, IP3 receptor and cGMP kinase Ibeta. Nature 404, 197–201.

Schlossmann, J., and Hofmann, F. (2005). cGMP-dependent protein kinases in drug discovery. Drug Discov Today 10, 627–634.

Serulle, Y., Zhang, S., Ninan, I., Puzzo, D., McCarthy, M., Khatri, L., Arancio, O., and Ziff, E.B. (2007). A GluR1-cGKII interaction regulates AMPA receptor trafficking. Neuron 56, 670–688.

Smith, J.A., Francis, S.H., Walsh, K.A., Kumar, S., and Corbin, J.D. (1996). Autophosphorylation of type Ibeta cGMP-dependent protein kinase increases basal catalytic activity and enhances allosteric activation by cGMP or cAMP. J Biol Chem 271, 20756–20762.

Surks, H.K., Mochizuki, N., Kasai, Y., Georgescu, S.P., Tang, K.M., Ito, M., Lincoln, T.M., and Mendelsohn, M.E. (1999). Regulation of myosin phosphatase by a specific interaction with cGMP-dependent protein kinase Ialpha. Science 286, 1583–1587.

Taylor, S.S., Kim, C., Cheng, C.Y., Brown, S.H., Wu, J., and Kannan, N. (2008). Signaling through cAMP and cAMP-dependent protein kinase: diverse strategies for drug design. Biochim Biophys Acta 1784, 16–26.

Taylor, S.S., Soberg, K., Kobori, E., Wu, J., Pautz, S., Herberg, F.W., and Skalhegg, B.S. (2021). The tails of PKA. Mol Pharmacol.

Vaandrager, A.B., Smolenski, A., Tilly, B.C., Houtsmuller, A.B., Ehlert, E.M., Bot, A.G., Edixhoven, M., Boomaars, W.E., Lohmann, S.M., and de Jonge, H.R. (1998). Membrane targeting of cGMP-dependent protein kinase is required for cystic fibrosis transmembrane conductance regulator Cl-channel activation. Proceedings of the National Academy of Sciences of the United States of America 95, 1466–1471.

VanSchouwen, B., Akimoto, M., Sayadi, M., Fogolari, F., and Melacini, G. (2015a). Role of Dynamics in the Autoinhibition and Activation of the Hyperpolarization-activated Cyclic Nucleotide-modulated (HCN) Ion Channels. J Biol Chem 290, 17642–17654.

VanSchouwen, B., and Melacini, G. (2018). Role of Dimers in the cAMP-Dependent Activation of Hyperpolarization-Activated Cyclic-Nucleotide-Modulated (HCN) Ion Channels. J Phys Chem B 122, 2177–2190.

VanSchouwen, B., Selvaratnam, R., Giri, R., Lorenz, R., Herberg, F.W., Kim, C., and Melacini, G. (2015b). Mechanism of cAMP Partial Agonism in Protein Kinase G (PKG). J Biol Chem 290, 28631–28641.

Wall, M.E., Francis, S.H., Corbin, J.D., Grimes, K., Richie-Jannetta, R., Kotera, J., Macdonald, B.A., Gibson, R.R., and Trewhella, J. (2003). Mechanisms associated with cGMP binding and activation of cGMP-dependent protein kinase. Proceedings of the National Academy of Sciences of the United States of America 100, 2380–2385.

## References

Cook, P. F., Neville, M. E., Vrana, K. E., Hartl, F. T., & Roskoski, R. (1982). Adenosine cyclic 3’,5’-monophosphate dependent protein kinase: kinetic mechanism for the bovine skeletal muscle catalytic subunit. Biochemistry, 21(23), 5794–5799. https://doi.org/10.1021/bi00266a011

